# Genetic signatures of evolutionary rescue by a selective sweep

**DOI:** 10.1101/800201

**Authors:** Matthew M. Osmond, Graham Coop

## Abstract

One of the most useful models in population genetics is that of a selective sweep and the consequent hitch-hiking of linked neutral alleles. While variations on this model typically assume constant population size, many instances of strong selection and rapid adaptation in nature may co-occur with complex demography. Here we extend the hitch-hiking model to evolutionary rescue, where adaptation and demography not only co-occur but are intimately entwined. Our results show how this feedback between demography and evolution determines – and restricts – the genetic signatures of evolutionary rescue, and how these differ from the signatures of sweeps in populations of constant size. In particular, we find rescue to harden sweeps from standing variance or new mutation (but not from migration), reduce genetic diversity both at the selected site and genome-wide, and increase the range of observed Tajima’s *D* values. For a given initial rate of population decline, the feedback between demography and evolution makes all of these differences more dramatic under weaker selection, where bottlenecks are prolonged. Nevertheless, it is likely difficult to infer the co-incident timing of the sweep and bottleneck from these simple signatures, never-mind a feedback between them. Temporal samples spanning contemporary rescue events may offer one way forward.

## Introduction

The simple models used to predict the genetic signatures of selective sweeps have been incredibly helpful in understanding and identifying population genetic signals of adaptation (e.g., Maynard Smith and Haigh, 1974; Kaplan et al., 1989; reviewed in Stephan, 2019). These models are usually based on constant-sized, Wright-Fisher populations. Meanwhile, many instances of adaptation – and thus selective sweeps – in nature will co-occur with complex demography. In fact, many of the most well-known examples of selective sweeps have arisen following a rather extreme and sudden change in the environment (e.g., after the application of insecticides, Sedghifar et al., 2016, or antimalarial drugs, Nair et al., 2003), which could have simultaneously imposed sharp demographic declines. Attempts to capture such complex demographic scenarios typically impose qualitatively appropriate changes in population size (e.g., Hermisson and Pennings, 2005). Indeed, a number of studies have explored the genetic signatures of selective sweeps during demographic bottlenecks (e.g., Innan and Kim, 2004; Teshima et al., 2006; Wilson et al., 2014). However, these demographies are nearly always chosen in the absence of an explicit population model and independently of evolution.

Here we model selective sweeps in a scenario where demography and adaptive evolution are not independent. In particular we model an instance of evolutionary rescue (Gomulkiewicz and Holt, 1995; reviewed in Bell, 2017), where a sudden environmental change causes population decline that is reverted by a selective sweep. Under this framework, rescue is a simultaneous demographic bottleneck and selective sweep, where each affects the other. First, because the mean absolute fitness of the population changes with the beneficia allele’s frequency, the depth and duration of the bottleneck depends on the dynamics of the selective sweep, i.e., evolution affects demography. Second, the probability that the beneficial allele establishes depends on its growth rate and thus the rate of population decline, i.e., demography affects evolution. Together, this feedback between demography and evolution restricts the range of dynamics that are possible, and therefore also restricts the range of genetic signatures we should expect to observe. Our goal here is to describe the range of genetic signatures that (this model of) evolutionary rescue allows, to help elucidate how rescue may obscure inferences of past selection and demography and to identify patterns that could be used to infer rescue in nature.

Most theory on evolutionary rescue to date (reviewed in Alexander et al., 2014) has focused on the probability of rescue. Recently, however, some attention has been given to the dynamics of population size (Orr and Unckless, 2014), the probability of soft sweeps (Wilson et al., 2017), and the genetic basis of adaptation (Osmond et al., 2020) *given* rescue in haploid or asexual populations. Here we extend this line of thinking to three modes of rescue in diploid, sexual populations, and use coalescent theory and simulations of whole chromosomes to examine the genetic signatures at linked, neutral loci. Our focus is on three common genetic signatures: the number of unique lineages of the beneficial allele that establish (i.e., the softness of the sweep), the pattern of nucleotide diversity around the selected site (i.e., the dip in diversity), and the pattern of Tajima’s *D* around the selected site (i.e., skews in the site-frequency spectrum). We explore three modes of rescue, where the beneficial allele arises from either standing genetic variance, recurrent *de novo* mutation, or migration, and compare to selective sweeps arising in populations of constant size. Qualitatively, we find that for a given selection coefficient rescue causes faster, harder sweeps that produce wider, deeper dips in diversity and more extreme values of Tajima’s *D*. Due to the feedback between demography and evolution, the effect of rescue on the signatures of selective sweeps becomes more pronounced as the selection coefficient, and thus the probability of rescue, gets smaller.

## Materials and Methods

### Population dynamics

#### Deterministic trajectories

Consider a population of size *N*(*t*) with a beneficial allele, *A*, at frequency *p*(*t*) and an ancestral allele, *a*, at frequency 1 − *p*(*t*). Assume non-overlapping generations and let *W*_*AA*_, *W*_*Aa*_, and *W*_*aa*_ be the absolute fitness (expected number of offspring) of each genotype. We are interested in the scenario where a population composed of primarily *aa* genotypes is declining at some rate *d*, i.e., *W*_*aa*_ = 1 − *d*. The beneficial allele, *A*, is then assumed to act multiplicatively with the fitness of the ancestral background, such that *W*_*Aa*_ = (1 − *d*)(1 + *hs*) and *W*_*AA*_ = (1 − *d*)(1 + *s*). Throughout the text we assume random mating, weak selection, *s* << 1, and additivity at the selected locus, *h* = 1/2, such that the allele frequency (c.f., equation 5.3.12 in Crow and Kimura, 1970) and population dynamics 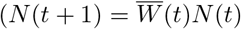, with 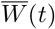 the population mean fitness) can be approximated by

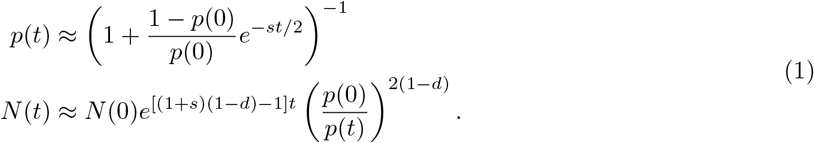

Equation 1 allows for unbounded population growth; to prevent this we impose a simple carrying capacity at *N*(0) by setting *N*(*t*) to be the minimum of *N*(0) and the value given in Equation 1.

#### Effective initial allele frequency

Equation 1 considers only the deterministic dynamics. We are, however, primarily concerned with a stochastic event: the establishment of a (potentially rare) beneficial allele. In rescue we are particularly interested in only those instances where the beneficial allele establishes – otherwise the population is extinct. Conditioning on establishment creates a bias away from the deterministic trajectory. As shown in Maynard Smith (1971) (see Orr and Unckless, 2014, for an application to evolutionary rescue), we can approximate this bias by assuming that a single copy of an allele that establishes with probability *P*_est_ instantaneously increases from an initial frequency of 1/(2*N*) to 1/(2*NP*_est_). In essence, dividing the initial allele frequency by the probability of establishment implies that an allele escaping random loss when rare will quickly increase to a larger frequency than otherwise expected before settling into its deterministic trajectory. As we will see below, we can combine this logic with the waiting time distribution for an establishing mutation to arrive to derive an effective initial allele frequency, *p*_0_. Using the deterministic prediction (Equation 1) with *p*(0) = *p*_0_ then approximates the forward-time trajectory of an allele that is destined to fixation.

We derive the probability of establishment, *P*_est_, assuming alleles do not interact (i.e., a branching process). Under this assumption a single copy of an allele that is expected to leave 1 + *ϵ* copies in the next generation with a variance of *v* has an establishment probability of (Allen, 2010, p. 172, see also Feller, 1951, equation 5.7, for a derivation from a diffusion approximation)

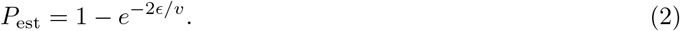

When *d* ≤ 0 we ignore density-dependence and the growth rate of the establishing heterozygote then depends only on its absolute fitness, *ϵ* = *W*_*Aa*_ − 1 = (1 + *sh*)(1 − *d*) − 1 ≈ *sh − d*. The variance, *v*, depends on particulars of the lifecycle (see Supplementary material: Genetic drift in the simulated lifecycle for details). When *h* = 1 and *v* = 1 we retrieve the probability of establishment of a weakly beneficial mutation in an exponentially growing or declining haploid population with a Poisson offspring distribution, *P*_est_ ≈ 2(*s − d*) (Otto and Whitlock, 1997).

#### Effective final allele frequency

Above we have stitched together the initial stochastic phase of the allele frequency and population dynamics with the deterministic phase by using *p*(0) = *p*_0_. We next incorporate the final phase, fixation, which is also stochastic. Incorporating this final phase is important here (as opposed to studies that predict only the forward-time dynamics; e.g., Orr and Unckless, 2014) because in looking at genetic signatures we are primarily concerned with the dynamics backwards in time from the point of fixation, and this third phase determines how fast fixation occurs.

As shown in Martin and Lambert (2015), we can treat the backward-in-time dynamics from fixation as we did the forward-in-time dynamics at establishment. In particular, the probability the heterozygote establishes (backward in time) in a population of mutant homozygotes is also given by Equation 2 but with the growth rate of a rare heterozygote in this population, *ϵ*, replaced by −*ϵ* (let’s call this probability 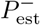). For instance, if the fitness of the heterozygote is *W*_*Aa*_ = (1 − *d*)(1 + *sh*) and the fitness of the mutant homozygote is *W*_*AA*_ = (1 − *d*)(1 + *s*) and the population is at carrying capacity then the growth rate of the heterozygote depends only on its relative fitness, −*ϵ* = 1 − *W*_*Aa*_/*W*_*AA*_ = 1 − (1 + *sh*)/(1 + *s*) ≈ *s*(1 − *h*), which is independent of demography, *d*.

Given the heterozygote establishes (backward-in-time) it will quickly increase to 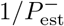 copies, implying that in a population of size *N* fixation effectively occurs when the deterministic trajectory reaches a frequency of 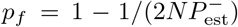). Below we will explore scenarios where fixation is expected to occur while the population is at carrying capacity, *N* (0). Thus in all cases we have the same effective final allele frequency,

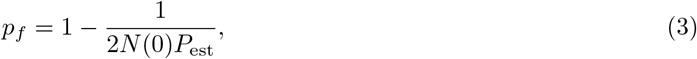

where *P*_est_ is given by Equation 2 with *ϵ* = 1 − (1 + *sh*)/(1 + *s*) and *v* as given in Supplementary material: Genetic drift in the simulated lifecycle.

#### Time to fixation

We can estimate the time to fixation in the additive model (*h* = 1/2) as the time it takes the deterministic approximation (Equation 1) to go from the effective initial frequency, *p*_0_, to the effective final frequency, *p*_*f*_, which is (equation 5.3.13 in Crow and Kimura, 1970)

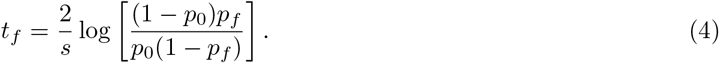

#### Backward-time dynamics

Setting *t* = *t*_*f*_ − *τ* in Equation 1 gives an approximation of the dynamics of allele frequency and population size backward in time,

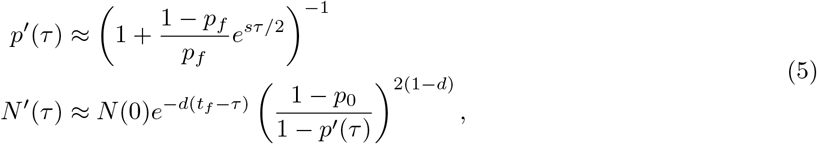

from the point of fixation, *τ* = 0, to the time of establishment, *τ* = *t*_*f*_. Note that the backward-time allele frequency dynamics do not depend on the initial frequency, *p*_0_, or demography, *d*, except in determining the maximum value of *τ*, *t*_*f*_. Thus the backward-time allele frequency dynamics are expected to be the same in rescue as in populations of constant size with the same carrying capacity (such that the final frequency, *p*_*f*_, is the same). Meanwhile, the backward-time population size dynamics depend heavily on the initial frequency as the time at which the sweep establishes determines the depth and duration of the bottleneck.

### The structured coalescent

#### Event rates

To explore the genetic patterns created by evolutionary rescue we next consider a random sample of chromosomes at the time the beneficial allele fixes. Focusing on a neutral locus that is recombination distance *r* from the selected site, we are interested in calculating the rate of coalescence, the rates of recombination and mutation off the selected background, and the rate of migration out of the population. If our sample of alleles has *k* distinct ancestors on the selected background *τ* generations before fixation, these rates are approximately (table 1 in Hudson and Kaplan, 1988; equation 16 in Pennings and Hermisson, 2006*a*; see Supplementary material: Deriving the structured coalescent for derivations)

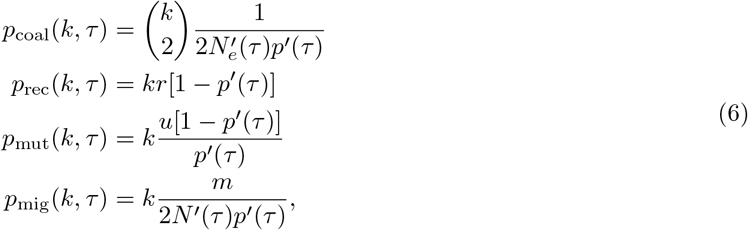

where 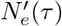 is the effective population size *τ* generations before fixation (see Supplementary material: Genetic drift in the simulated lifecycle for details).

#### Event timing

We now use the instantaneous event rates (Equation 6) to calculate the probability that the most recent event is *τ* generations before fixation and is either coalescence, recombination, mutation, or migration. Letting *i, j* ∈ {coal, rec, mut, mig}, the probability that *i* is the most recent event and occurs *τ* generations before fixation is (c.f., equation 6 in Pennings and Hermisson, 2006*b*)

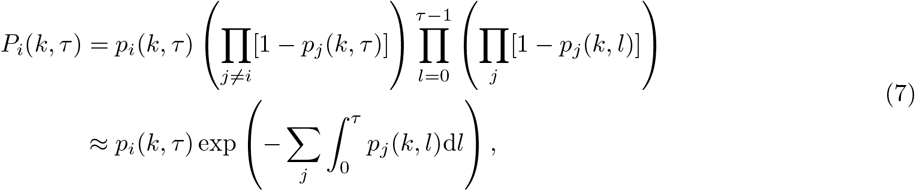

i.e., the waiting time for an inhomogeneous exponential random variable. The approximation assumes the *p*_*i*_(*k, τ*) are small enough such that at most one event happens each generation, with small probability, and the changes in the *p*_*i*_(*k, τ*) from one generation to the next are small enough that we can approximate a sum across *τ* with an integral. As a technical aside, to speed computation we analytically solve the integrals in Equation 7 under the assumption of unbounded population growth (Equation 1).

### Genetic signatures at linked neutral loci

#### Pairwise diversity

One classic signature of a selective sweep is a dip in genetic diversity around the selected site (Maynard Smith and Haigh, 1974; Kaplan et al., 1989). Here we consider the average number of nucleotide differences between two randomly sampled sequences, *π* (Tajima, 1983), focusing on sequences of length 1 (i.e., heterozygosity).

We first consider our expectation for *π* at a site that is far enough away from the selected site to be unaffected by hitchhiking, which will provide us with an expectation for genome-wide average diversity. While not directly impacted by the sweep, these loci are indirectly impacted as the sweep dictates the severity of the population bottleneck. Our expectation for *π* at such a site in a population of constant effective size, *N*_*e*_, is simply 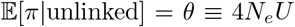 (Watterson, 1975), with *U* the per base per generation mutation rate at neutral loci. Ignoring neutral mutation input during the sweep, the *π* at a sufficiently loosely linked site in a population of changing size is this neutral expectation times the probability a sample of size two does not coalesce during the sweep (c.f., equation 4 in Slatkin and Hudson, 1991 and equation 7 in Griffiths and Tavare, 1994),

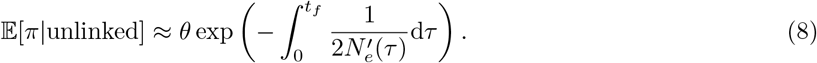

We next consider sites that are close enough to the selected locus that they are directly affected by the selective sweep through hitchhiking. To keep the analysis simple, we assume that if recombination or mutation moves one of the sampled alleles to the ancestral background before the two samples coalesce then it is as if both samples were on the ancestral background from the start and therefore coalesce with each other as if they were at an unlinked locus (Equation 8). This assumption ignores the time it takes for the two samples to arrive on the ancestral background and is therefore expected to underestimate diversity at moderately-linked sites. There is also the possibility that no events occur in the history of the sample during the sweep. Then, if the sweep arose from mutation or migration or from a single copy of the beneficial allele (*κ* = 1), the sample must coalesce. If we instead start with more than one beneficial copy, *κ* > 1, it is possible that the two samples had ancestors that were linked to distinct copies of the beneficial allele within the standing variation. While the coalescent naturally tells us the probability that two lineages do not coalesce during the sweep (since distinct copies of the beneficial allele are exchangeable; e.g., Tavaré, 1984), our continuous-time approximation of the coalescent (Equation 7) implicitly assumes a large number of copies of the beneficial allele at any one time. To account for the fact that *κ* is finite, and often small, we assume that if under the continuous-time approximation of the coalescent there remain two lineages on the beneficial background at the beginning of the sweep then they coalesce with probability 1/*κ* (see the ‘instantaneous’ approximation in Anderson and Slatkin, 2007, for a similar approach). Finally, we assume that each copy of the beneficial allele in the standing variance has a recent and unique mutational origin (implying the beneficial allele was sufficiently deleterious before the environmental change; c.f., Prezeworski et al., 2005; Berg and Coop, 2015), so that two distinct ancestral lineages are independent draws from a neutral population. Ignoring neutral mutations during the sweep, a simple approximation for diversity at any location in the genome is then

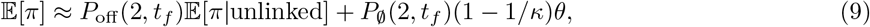

where 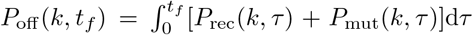 is the probability of recombination or mutation befor ecoalescence during the sweep and 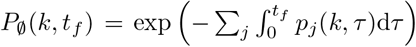 is the probability that no events have occurred in the history of the sample during the sweep (*j* ∈ {rec, mut, coal}). Equation 9 is a function of recombination distance, *r*, and approaches 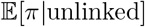 as *r* gets large.

#### Tajima’s *D*

As a second genetic signature at linked neutral sites we consider Tajima’s *D* statistic (Tajima, 1989), which measures the excess (positive *D*) or deficiency (negative *D*) of intermediate frequency polymorphisms relative to the standard neutral model. Quantitative predictions of Tajima’s *D* require one to consider samples of size greater than 2, which quickly becomes complicated with selection and non-equilibrial demography. Instead, here we discuss the expected qualitative patterns based on intuition from the analysis presented above.

First, hard selective sweeps tend to produce star-like gene genealogies, with most samples coalescing near the beginning of the sweep and recombination allowing a few samples to coalesce much further back in time (Kaplan et al., 1989). Hard sweeps therefore produce an excess of low and high frequency polymorphisms (Wakeley, 2009, p. 120), leading to negative *D* (Braverman et al., 1995). Larger selection coefficients create more star-like the genealogies (as there is then less time for recombination off the sweep) and thus more negative *D* when conditioned on a hard sweep. However, with sufficient standing genetic variation or rates of recurrent mutation or migration, larger selection coefficients will tend to cause softer selective sweeps as they increase the probability any one copy establishes (Equation 2). Soft selective sweeps (by definition) allow some samples to coalesce further back in time, before the start of the sweep, even at the selected site. Soft sweeps therefore tend to have less effect on neutral genealogies and hence on *D*, although sufficiently soft sweeps can actually cause positive *D*, by allowing intermediate-sized groups of samples to descend from different ancestors containing the beneficial allele (Pennings and Hermisson, 2006*b*).

As linkage to the selected site decreases so too does this skew in genealogies. In the case of a constant population size, *D* should asymptote to the neutral expectation of zero. In the case of rescue, however, the bottleneck will cause an excess of intermediate frequency polymorphisms (Wakeley, 2009, p. 120), and therefore *D* should asymptote at some positive value (more positive with more severe bottlenecks).

### Simulations

The simulation details are described in full in Supplementary material: Simulation details. Briefly, the lifecycle described in Supplementary material: Simulated lifecycle was simulated in `SLiM 3` (Haller and Messer, 2019) with tree-sequence recording (Haller et al., 2019). We simulated a 20 Mb segment of a chromosome, with all but one of the center loci neutral, with a per base recombination rate of *r*_*bp*_ = 2 × 10^−8^ and per base mutation rate at neutral loci of *U* = 6 × 10^−9^ (both inspired by *Drosophila* estimates; Mackay et al., 2012; Haag-Liautard et al., 2007). A simulation was considered successful and ended when the beneficial mutation was fixed and the population size had recovered to its initial size, *N*(0). Successful simulations were recapitated with Hudson’s coalescent algorithm (Hudson, 1983, 2002), implemented in `msprime` (Kelleher et al., 2016), using *N*_*e*_ ≈ 4*N*(0)/7 (see Supplementary material: Genetic drift in the simulated lifecycle). Pairwise diversity, Tajima’s *D*, site frequency spectra, and linkage disequilibrium were then calculated directly from the tree sequences using `tskit` (Kelleher et al., 2018), for a random sample of 100 contemporary chromosomes. We chose to simulate a population with initial census size *N*(0) = 10^4^ declining at rate *d* = 0.05, where the beneficial allele had selection coefficient *s* = 0.13 or *s* = 0.20. This describes a relatively small population that is expected to go extinct in ≈ 200 generations in the absence ofa sweep at a locus under strong selection (as might be expected following a severe environmental change).

### Data availability statement

Code used to derive and plot all results (Python scripts and Mathematica notebook) is available at https://github.com/mmosmond/rescueCoalescent.git. File S1 refers to the Mathematica notebook, which is also provided in freely accessible forms (e.g., PDF).

## Results

### Rescue from standing genetic variation (SGV)

We first consider rescue arising from genetic variation that is present at the time of the environmental change and ignore further mutations and migration. To avoid complicating the presentation we assume the initial number of beneficial alleles is given; treating the initial number as a random variable requires conditioning the distribution on a successful sweep (Hermisson and Pennings, 2017).

#### The probability of rescue and soft selective sweeps

Given there are initially *κ* « *N*(0) copies of the beneficial allele, the number that establish is roughly binomially distributed with *κ* trials and success probability *P*_est_. The probability of rescue is the probability that at least one establishes (c.f., equation 2 in Orr and Unckless, 2014),

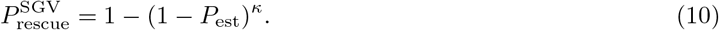

Conditioning on rescue, the expected number of establishing copies is

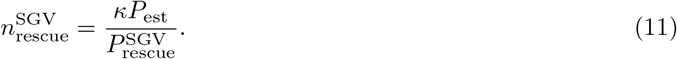

Note that when the unconditioned expected number of establishing copies, *κP*_est_, is small, the probability of rescue is roughly *κP*_est_. Equation 11 then implies we expect only one of the initial copies to establish, i.e., we expect a hard sweep. For larger values of *κP*_est_ rescue will often occur by a soft selective sweep (Hermisson and Pennings, 2005), where multiple initial copies establish. The probability that multiple copies establish is the probability of rescue minus the probability of a hard sweep, 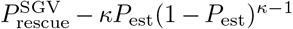. Given rescue occurs, the probability it is due to a soft sweep can therefore be written as

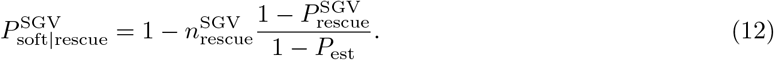

As expected, the probability of a soft sweep given rescue is 0 when we start with a single copy, *κ* = 1, and approaches 1 as the number of copies becomes large. Between these two extremes we find that Equation 12 provides reasonable estimates for small *d* or large *s* but underestimates the probability of a soft sweep otherwise (Figure 1A), when beneficial alleles can persist at low numbers long enough to establish with some non-negligible probability of experiencing some selection as homozygotes (thus increasing their probability of establishment). Using the probability of establishment observed in the simulations corrects this error (see File S1). Because the expected number of copies that establishes given rescue (Equation 11) is nearly independent of *P*_est_ (as long as it is small), Equation 11 more closely matches simulations (Figure 1B).

**Figure 1:**
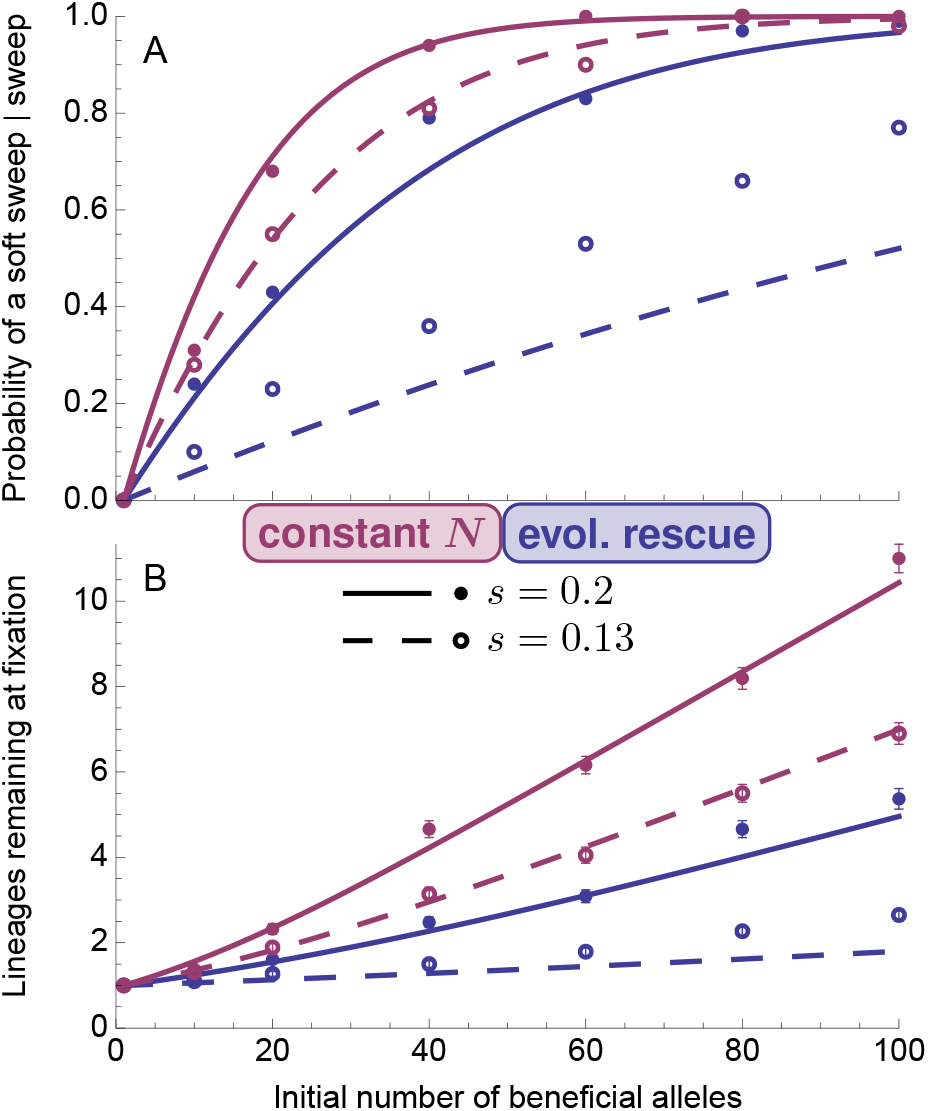
Soft sweeps from standing genetic variation. (A) The probability that more than one initial copy of the beneficial allele has descendants at the time of fixation in rescue (blue; *d* = 0.05) or in a population of roughly constant size (red; *d* = 0) as a function of the initial number of copies of the beneficial allele, *κ*. (B) The expected number of initial copies of the beneficial allele that have descendants at the time of fixation. The curves are Equations 12 (panel A) and 11 (panel B). Each point is based on 100 replicate simulations where a sweep (and population recovery) was observed. Error bars in (B) are standard errors. Parameters: *N*(0) = 10^4^.

#### Effective initial allele frequency and the backward-time dynamics

As each of the 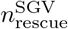 expected establishing copies given rescue is expected to rapidly reach 1*/P*_est_ copies (see Effective initial allele frequency), it is as if the sweep began deterministically (Equation 1) with initial frequency (c.f., equation S1.4 in Orr and Unckless, 2014)

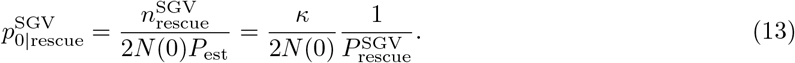

This same conditioning applies in a population of constant size (i.e., *d* = 0). As the decline rate, *d*, increases the probability a copy establishes, *P*_est_, and hence the probability of rescue, 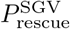, declines, making the conditioning stronger (i.e., increasing the difference between the effective and true initial allele frequencies). This causes selective sweeps to get started faster as *d* increases, implying that, all else equal, rescue sweeps are expected to be shorter than those in populations of constant size.

Using 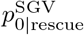 as *p*_0_ in Equations 4 and 5 gives our semi-deterministic prediction of the backward-time dynamics. As shown in Figure 2, this does a reasonably good job of describing the simulation results.Looking closer, we find that our predictions do an excellent job of approximating the mean allele frequency as it departs from 1, but typically begin to underestimate the mean allele frequency as frequencies drop lower (see File S1 for more detail). Consequently, we tend to underestimate the mean time to fixation (compare arrows and stars in Figure 2) and overestimate mean population size. This error arises even under a true branching process (e.g., under clonal reproduction) in a population of constant size and becomes worse as selection becomes weaker (see File S1). The reason for this error is that one minus the effective final allele frequency, 1 − *p*_*f*_ = *q*_*f*_ is exponentially distributed with expectation 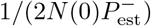 (Martin and Lambert, 2015), where 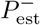 is the backward-time establishment probability of the heterozygote in a population of mutant homozygotes and we have assumed fixation occurs when the population size is *N*(0) (which is expected to be true in all of our numerical examples). The variance in the effective final allele frequencytherefore increases like 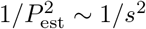 as *s* declines, meaning that under weaker selection there is much more variance in the time spent in the stochastic phase near fixation. Combining this distribution of waiting times to leave the stochastic phase with the non-linearities of the subsequent deterministic phase causes our semi-deterministic approximation to generally underestimate allele frequency. Despite this error we forge ahead and use our simple semi-deterministic approximations in the structured coalescent (Equations 6–7).

**Figure 2:**
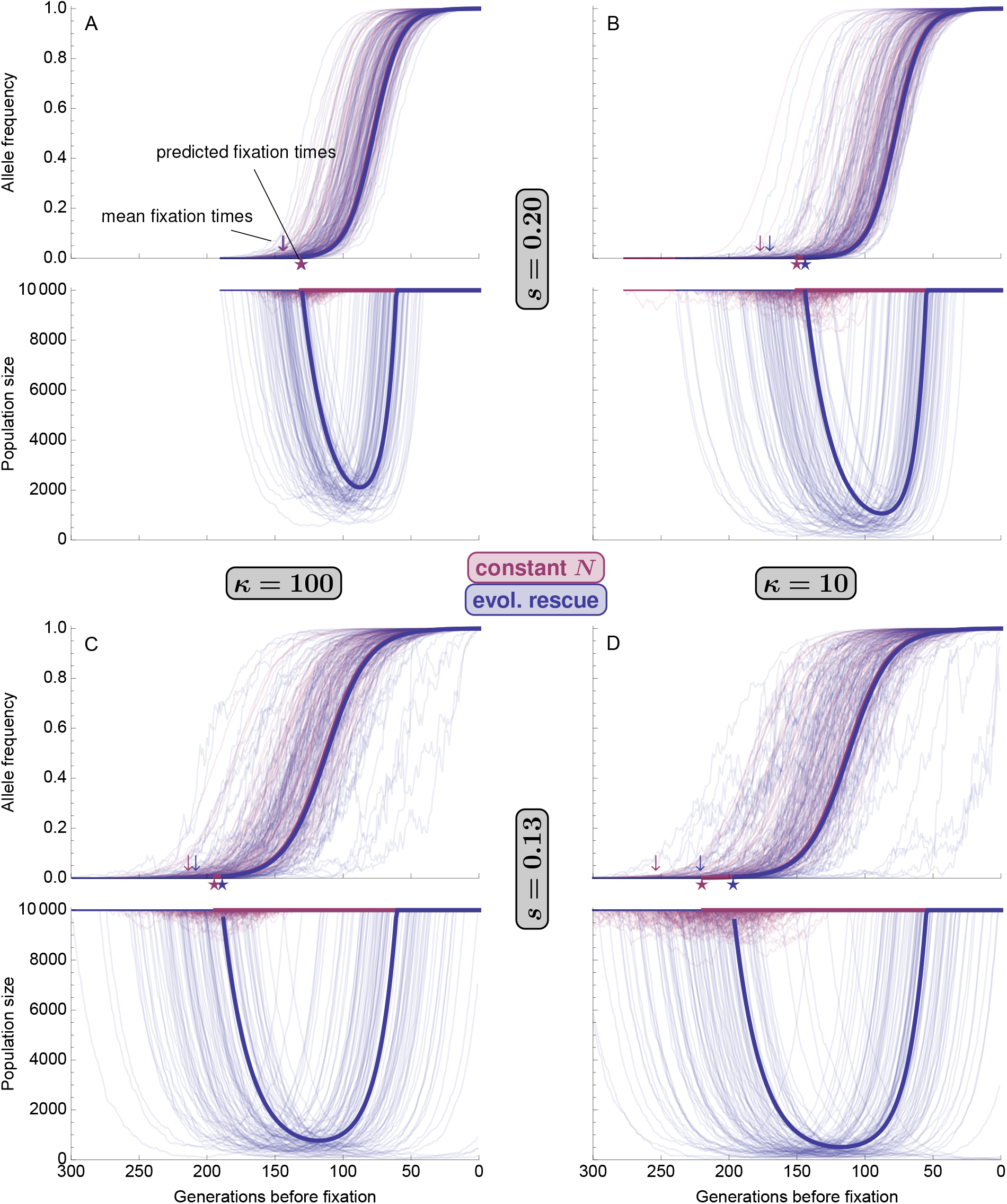
Allele frequency, *p*′(*τ*), and population size, *N*′(*τ*), at generation *τ* before fixation during a selective sweep from standing genetic variation in evolutionary rescue (blue; *d* = 0.05) and in a population of roughly constant size (red; *d* = 0). The thick solid curves are analytic approximations (Equation 5), using 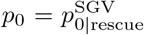 Equation 13) as the initial frequency and *p*_*f*_ (Equation 3) as the final allele frequency. The thin curves are 100 replicate simulations where a sweep (and population recovery) was observed. The stars show the predicted time to fixation (Equation 4). The arrows show the mean time to fixation observed in simulations.

Before moving on to the coalescent, however, a number of biological insights can be gleaned from Figure 2 that will help explain downstream results: 1) the backward-time allele frequency dynamics are the same in rescue as they are in populations of constant size since they only depend on the effective final allele frequency, *p*_*f*_, which does not depend on *d* (Equation 3), 2) fixation times are shorter in rescue as, forward-in-time, those sweeps tend to get started faster (i.e., the effective initial frequency increases with *d*; Equation 13), 3) weaker selection causes larger population bottlenecks as the deterministic portion of the sweep is then slower and a given frequency change has less impact on population mean fitness (Equation 1), and 4) both the allele frequency and population size dynamics depend only weakly on the initial number of mutants, despite large differences in the probability of rescue, because sweeps that are less likely to occur get started faster (Equation 13).

#### The structured coalescent

For a linked neutral allele in a sample of size 2, Figure 3 compares the predicted probability of recombination off the selective sweep (opaque dashed curves) and coalescence on the beneficial background (opaque solid curves) with the mean probabilities given the allele frequency and population size dynamics observed in simulations (transparent curves). We find that our approximations do relatively well near fixation, where our predictions of allele frequency are better (Figure 2), but that our underestimates of allele frequency at later times cause us to generally overestimate rates of recombination and coalescence on the beneficial background near the beginning of the sweep. Despite this error we capture the qualitative dynamics and can draw out a number of interesting biological consequences: 1) the timing of the bottleneck during rescue pushes coalescent times towards the present, so that the distributions of coalescence and recombination times overlap more than in populations of constant size, 2) the bottleneck also increases the overall probability of coalescence in rescue, which reduces the probability of recombination off the sweep (compare areas under dashed curves), and 3) the difference in the structured coalescent between rescue and a population of constant size is larger under weaker selection. This latter point nicely illustrates the coupling between demography and evolution in rescue; while weaker selection creates a slower sweep and hence more time for recombination off the beneficial background, it also slows population recovery, leading to longer and deeper bottlenecks that increase coalescence enough to counteract the additional time provided for recombination.

**Figure 3:**
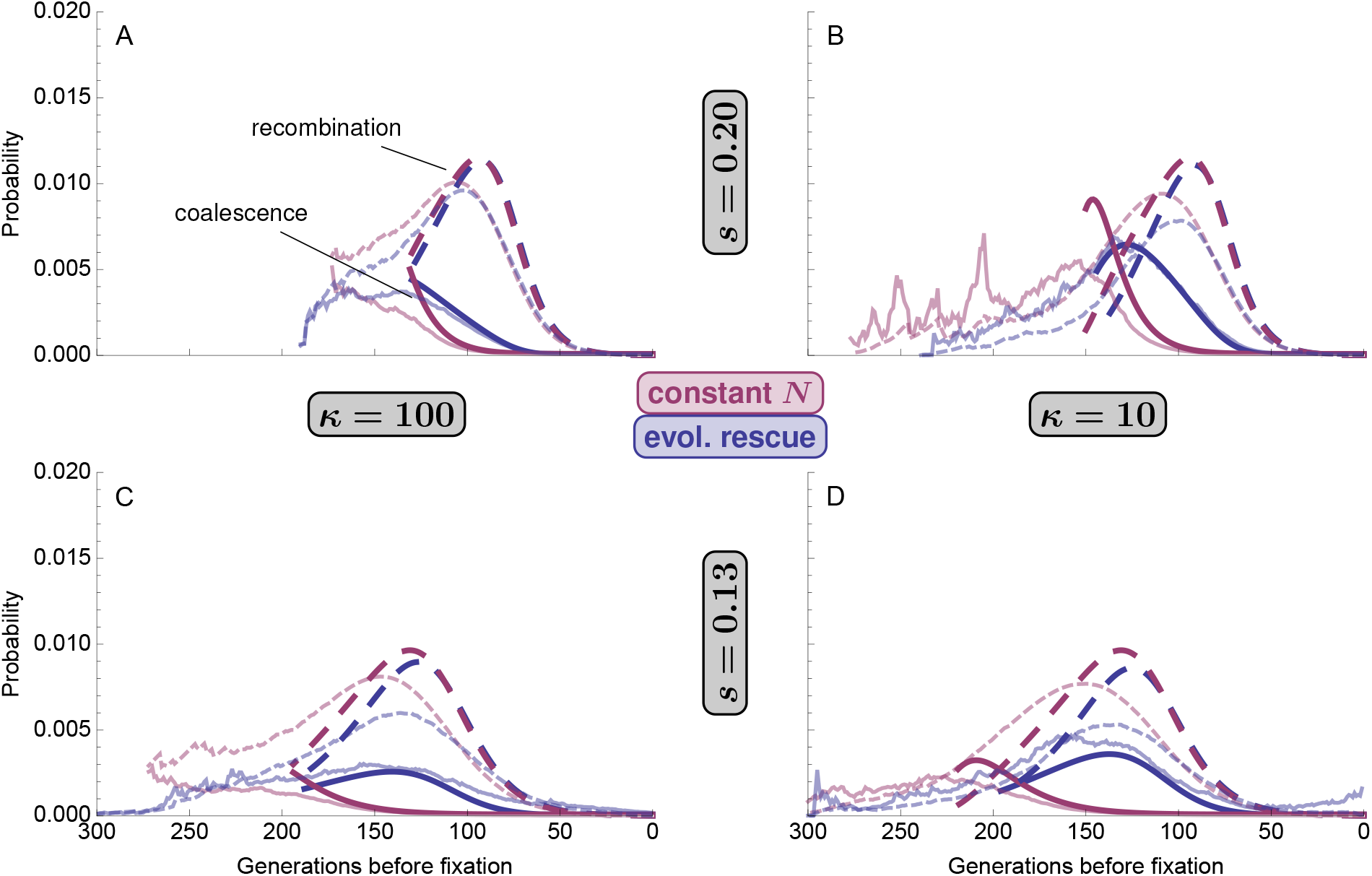
The timing of events in the structured coalescent (Equation 7) for a sample of size *k* = 2 at a linked neutral locus (*r* = 0.01) during a selective sweep from standing genetic variation in evolutionary rescue (blue; *d* = 0.05) or in a population of roughly constant size (red; *d* = 0). The opaque curves use the analytic expressions for allele frequency and population size (Equation 5) while the transparent curves show the mean probabilities given the observed allele frequency and population size dynamics in 100 replicate simulations (these become more variable back in time as less replicates remain polymorphic).

#### Genetic diversity

Figure 4 compares our predictions of relative pairwise diversity, 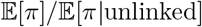 (Equations 8–9), against simulations, 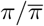. Despite the errors in the predictions of the dynamics above (Figure 2–3) our approximations capture the relative rates of recombination and coalescence. Two main conclusions emerge: 1) rescue tends to deepen dips in diversity when soft sweeps are possible, i.e., it hardens soft sweeps because the bottleneck increases the probability of coalescence (decreasing *P*_*∅*_(2, *T*) at the selected site, *r* = 0; Equation 9), and 2) rescue generally produces wider dips in diversity due to excess, and earlier, coalescence as well shorter sweeps (the magnitude and timing of the bottleneck, as well as lower establishment probabilities, decreases *P*_off_ (2, *T*) at a given *r* > 0; Equation 9). Note that when the probability of rescue is very small (Figure 4D), we predict the diversity at the selected site to actually be higher under rescue than in a population of constant size. This is driven by our prediction of fixation time; low probabilities of rescue imply high effective initial allele frequencies, leaving less time for coalescence to occur (compare solid curves in Figure 3D). This is prediction is not, however, borne out in the observed simulations, as we tend to overestimate coalescence in populations of constant size case and underestimate it in rescue (Figure 3D).

**Figure 4:**
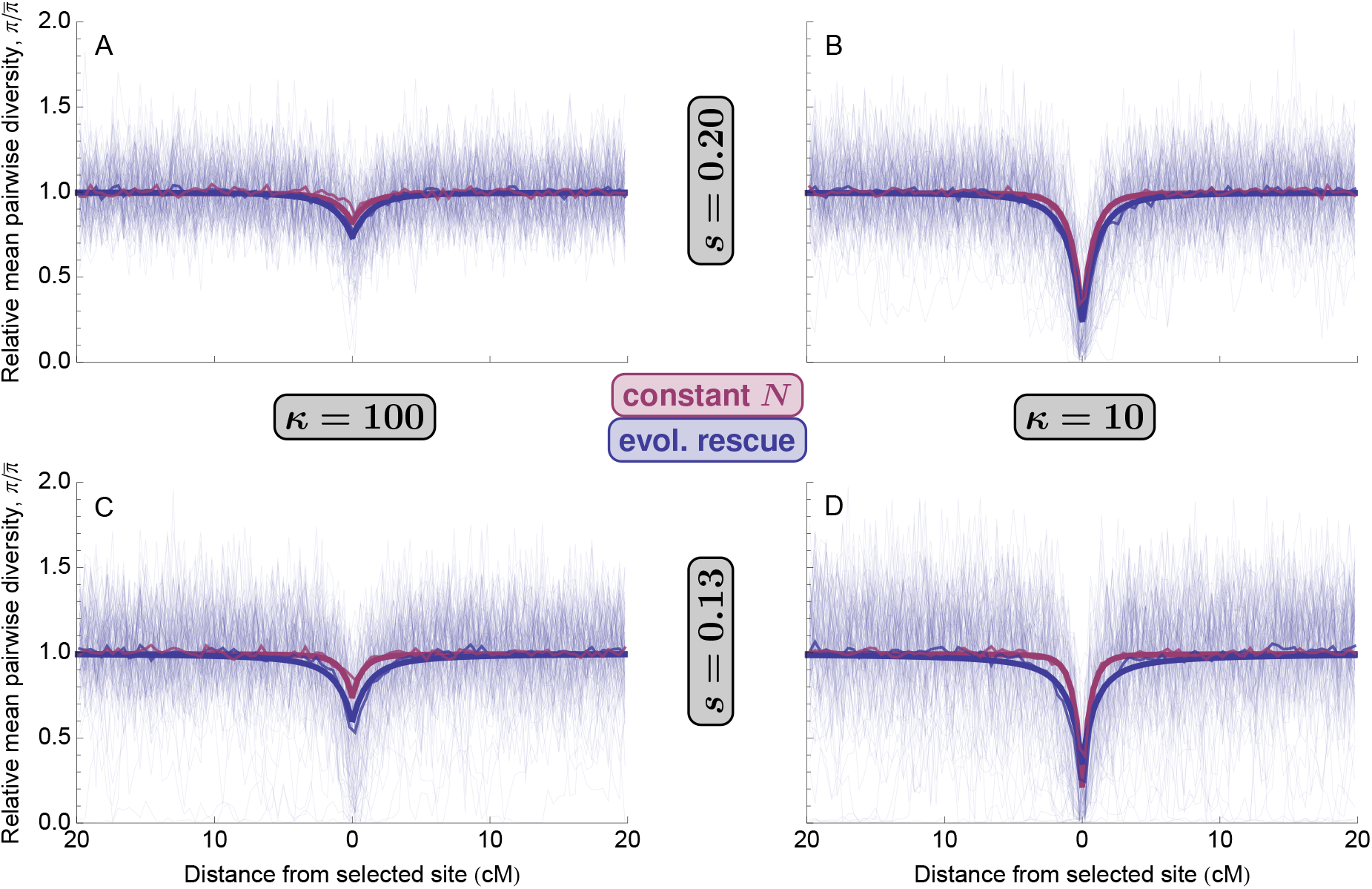
Relative mean pairwise diversity, 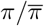, after a selective sweep from standing genetic variation during evolutionary rescue (blue; *d* = 0.05) or in a population of roughly constant size (red; *d* = 0). The thickest curves are 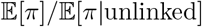 (Equation 9). The thinnest curves are 100 replicate simulations (rescue only for clarity) and the slightly thicker curves are simulation means (often obscured by prediction). Parameters: *N*(0) = 10^4^.

If the population was sampled both before and after the selective sweep (or we have very good estimates of its mutation rate and long-term effective population size), the absolute amount of pairwise diversity contains additional information. Figure S1 compares our predictions of 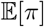 (Equation 9) against simulations, showing that the bottleneck decreases background levels of diversity as well. This decreases is more evident under weaker selection, where bottlenecks are more pronounced. While our predictions qualitatively match the mean simulation results, our tendency to underestimate allele frequencies and thus overestimate harmonic mean population sizes during rescue (Figures 2) causes us to generally overestimate background diversity in these cases. To correct for this, we can replace our prediction for background diversity, 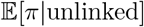 (Equation 8), in the rescue scenario with the observed genome-wide average diversity (dashed curves in Figure S1). A very similar result is achieved by using effective population sizes observed in simulations in Equation 8 (results not shown). Using the observed mean diversity level may be justified by the fact that genome-wide diversity can be measured directly from data and is highly variable across populations (Tajima, 1983).

#### Tajima’s *D*

Figure 5 shows the Tajima’s *D* values observed in simulations. There are two main take-aways: 1) the bottleneck during rescue causes positive background *D*, and 2) rescue sweeps tend to be harder and thus cause a greater decrease (or smaller increase) of *D* at the selected site. Together these patterns cause rescue to “stretch out” (e.g., panel D) or even invert (e.g., panel C) the pattern of *D* observed in populations of constant size.

**Figure 5:**
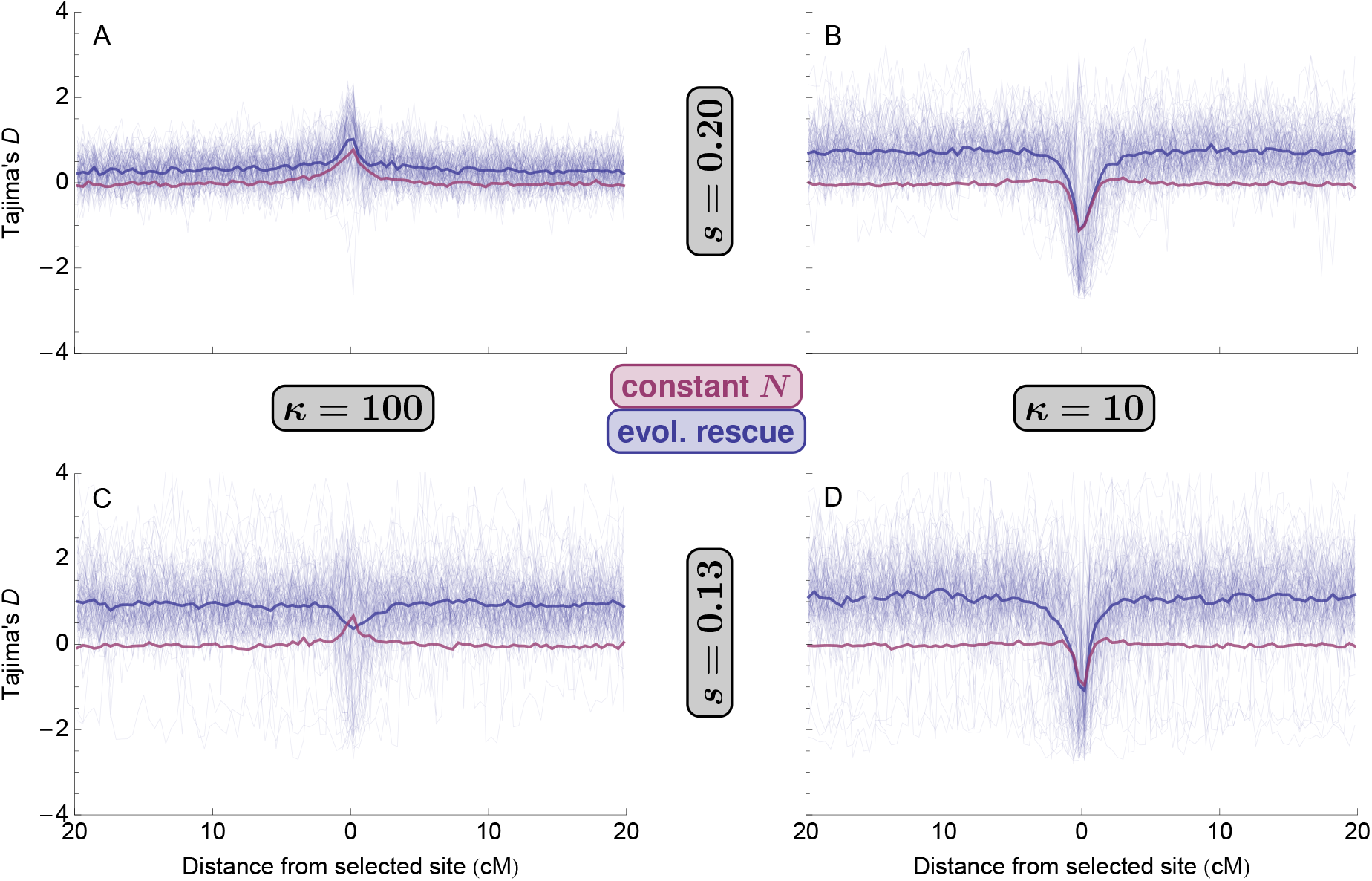
Tajima’s *D* after a selective sweep from standing genetic variation in evolutionary rescue (blue; *d* = 0.05) or in a population of roughly constant size (red; *d* = 0). Thin curves show 100 replicate simulations (rescue only for clarity) and thicker curves show simulation means. Parameters: *N*(0) = 10^4^.

### A simultaneous, but independent, sweep and bottleneck

It will of course be much harder to differentiate rescue from a simultaneous, but independent, sweep and bottleneck. In Supplementary material: A simultaneous, but independent, bottleneck and sweep from SGV we compare rescue, with expected effective population size *N*_*r*_ (from Equation 1), to sweeps that occur in populations suddenly bottlenecked to size *N*_*r*_. Because rescue has higher population sizes at the beginning and end of the sweep the sweeps take longer, leaving more time for coalescence, which leads to harder sweeps and lower genome-wide diversity. To see if the slightly different timings of recombination relative to coalescence are detectable, we examine the full site frequency spectrum as well as linkage disequilibrium. There is perhaps a slightly more uniform spectrum across intermediate frequency mutations at tightly linked sites under rescue, as expected given coalescence then overlaps more with recombination (similar to the effect of recurrent mutation Pennings and Hermisson, 2006*b*). Linkage disequilibrium is elevated under rescue, largely mirroring the patterns we see in absolute diversity.

### Rescue by *de novo* mutation (DNM)

#### The probability of rescue and soft selective sweeps

When there are few copies of the beneficial allele at the time of environmental change rescue may depend on mutations arising *de novo* at the selected site during population decline. To predict the allele frequency and population size dynamics in this scenario we then need to derive the waiting time until a rescue mutation successfully establishes. Ignoring unsuccessful mutations, the first successful rescue mutation arrives according to a time-inhomogeneous Poisson process with rate, *λ*(*t*) = 2*N*(*t*)*uP*^est^, where 2*N*(*t*) = 2*N*(0)*e*^−*dt*^ describes the decline in the number of copies of the ancestral allele. Thus the probability that a rescue mutation has established by time *T* is 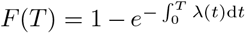. Taking the limit as time goes to infinity then gives the probability of rescue (c.f., equation 10 in Orr and Unckless, 2008)

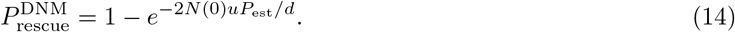

We can also calculate the probability of a soft sweep during rescue from recurrent mutation. Taking into account the beneficial alleles present at time *t*, the rate at which additional copies arise and establish is *λ*(*t*) = 2*N*(*t*)_*q*_(*t*)*uP*_est_(*t*), where 2*N*(*t*)_*q*_(*t*) is the number of ancestral alleles at time *t* and *P*est(*t*) is the probability of establishment at time *t*, which changes with allele frequency, and thus time, because allele frequency influences the genotypes the new allele experiences (i.e., its marginal fitness). Thus the number of mutations that arise and fix is Poisson with rate 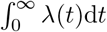. To gain intuition we make a very rough approximation, assuming *q*(*t*) ≈ 1 while mutations are arriving (i.e., while *N*(*t*) is still large), so that *P*_est_(*t*) ≈ *P*_est_ and we get the same Poisson rate we derived above for the first successful mutation, 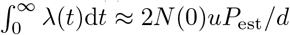. The resulting probability distribution for the number of mutations that establish is analogous to the result in a model with haploid selection (c.f., equation 7 in Wilson et al., 2017). Dividing the expected number of establishing mutations by the probability of rescue, the expected number that establish given rescue is

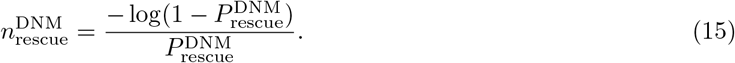

With these same approximations the probability of a soft selective sweep given rescue (i.e., the probability more than one copy establishes) is

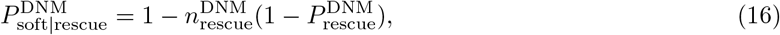

as in a haploid population (equation 8 in Wilson et al., 2017).

Both of these approximations tends to be underestimates (Figure 6), especially when selection is weak. This is partly because we also tend to underestimate the probability of rescue, which is elevated by increases in the frequency of the beneficial allele, which increases the marginal fitness of the ancestral allele whenever *h* > 0 (prolonging its persistence and therefore creating more opportunity for mutation) and increases the marginal fitness of the mutant allele (increasing the establishment probability). However, using the observed probability of rescue in Equations 15 and 16 still results in underestimates (see File S1) because ignoring increases in the beneficial allele frequency becomes less reasonable as more copies are established.

**Figure 6:**
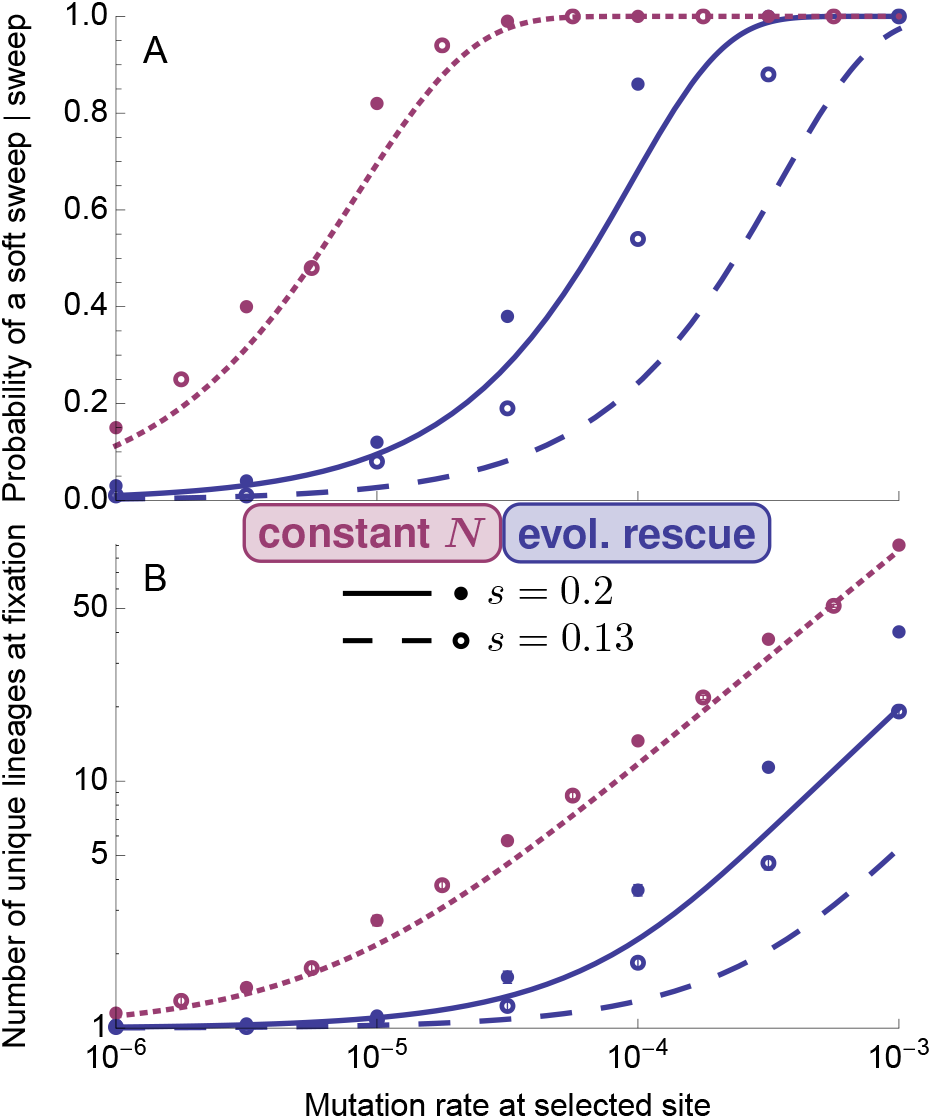
Soft sweeps from *de novo* mutation. (A) The probability that more than mutant lineage has descendants at the time of fixation in rescue (blue; *d* = 0.05) or in a population of roughly constant size (red; *d* = 0) as a function of the mutation rate, *u*. (B) The expected number of mutant lineages that have descendants at the time of fixation. The blue curves are Equations 16 (panel A) and 15 (panel B). The red curves are from Ewens’ sampling formula (equation 11 (panel A) and equation 12 (panel B) in Pennings and Hermisson, 2006*a*, with *θ* = 2*N*_*e*_(0)*u* = 8*N*(0)*u*/7 and *n* = 2*N*(0)). Each point is based on 100 replicate simulations where a sweep (and population recovery) was observed. Error bars in (B) are standard errors. Parameters: *N*(0) = 10^4^.

### Effective initial allele frequency and the backward-time dynamics

Following Orr and Unckless (2014) (see their Text S2 and our File S1 for more detail), taking the derivative of the cumulative distribution of waiting times for the first successful mutation, *F*(*T*), and dividing by the probability of rescue gives the probability distribution function for the arrival time of the first establishing rescue mutation given rescue, *f*(*t*). As discussed above, while the first establishing mutation is still rare it will grow exponentially at rate *ϵ* = (1+*sh*)(1−*d*)−1, and conditioned on its establishment will very quickly reach 1/*P*_est_ copies. Integrating over the distribution of arrival times then gives the expected number of copies of this successful mutation at time *t* since the environmental change given rescue, 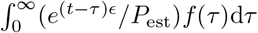. Solving this integral, evaluating at *t* = 0, and dividing by the total number of alleles at the time of environmental change, 2*N*(0), it is therefore as if the successful mutation was present at the time of environmental change, with frequency

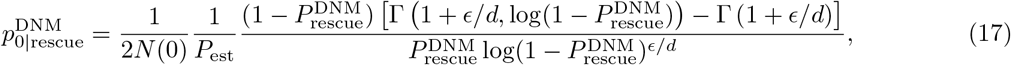

where Γ(*z*) is the gamma function (equation 6.1.1 in Abramowitz and Stegun, 1972) and Γ(*a, x*) is the incomplete gamma function (equation 6.5.3 in Abramowitz and Stegun, 1972). The factor 1/*P*_est_ > 1 increases the effective initial frequency, because we have conditioned on establishment, while the last factor decreases the effective initial frequency, because we must wait for the mutation to arise. When the unconditioned expected number of rescue mutations, 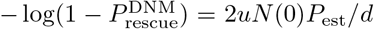, is small this cancels out and the last factor in Equation 17 becomes approximately *d*/(*d* + *ϵ*), which is independent of the mutation rate (as in the haploid case; Orr and Unckless, 2014). That is, conditioning on unlikely rescue, rescue mutations arise earlier in populations that decline faster.

With constant population size the time of establishment of the first successful mutation is irrelevant for the backward-in-time dynamics and therefore for the resulting genetic signatures (in contrast, when *d* > 0 we need to predict the time of establishment to predict the population size dynamics; Equation 5). For populations of constant size we need only determine the effective allele frequency at the time the sweep begins, which is simply 1/(2*N*(0)*P*_est_), independent of the mutation rate.

Figure S2 compares our analytical approximations of the dynamics (using Equations 5 and 17) against individual-based simulations. As with rescue from standing genetic variance (Figure 2), the non-linearities of the deterministic phase cause our semi-deterministic approximation to underestimate mean allele frequencies and overestimate mean population sizes as the allele frequency declines from fixation. We again see that rescue sweeps tend to be shorter than those in populations of constant size (due to lower establishment probabilities) and that the bottleneck sizes and sweep times given rescue depend little on the probability of rescue.

#### The structured coalescent

Figure S3 shows the timing of coalescence, recombination, and mutation for a sample of size 2 at a linked neutral locus. The patterns of coalescence and recombination are essentially the same as observed during sweeps from standing genetic variation (Figure 3). The timing of mutation is qualitatively similar to that of coalescence, peaking near establishment (since both depend on the inverse of allele frequency). The main effect of the excess coalescence during rescue is a reduced probability of mutating off the sweep (compare areas under dotted curves), which is exacerbated by rescue sweeps being shorter.

#### Genetic signatures at linked neutral loci

Figure S4 shows relative pairwise diversity around the selected site. The patterns here are nearly identical to those found after sweeps from standing genetic variation (Figure 4), although the difference between rescue and the constant population size model are larger here, perhaps partly due to the reduction in rates of mutation off the selected sweep (and also because our parameter choice causes the bottlenecks to be deeper when rescue occurs by mutation). Figure S5 shows Tajima’s *D* values around the selected site, which are nearly identical to those observed after sweeps from standing genetic variation (Figure 5).

On a related note, Wilson et al. (2017) reasoned that, because soft sweeps from recurrent mutation are expected when rescue is likely while hard sweeps are expected when rescue is rare (Equation 16), population bottlenecks will tend to be more extreme when rescue occurs by a hard selective sweep. From this they argued it might actually be easier to detect soft sweeps from patterns at linked neutral loci, as bottlenecks are expected to obscure the signal. Here we show the importance of conditioning on rescue, which roughly equalizes bottleneck sizes across scenarios with very different probabilities of rescue (e.g., due to differences in mutation rate; Figure S2), potentially making harder sweeps easier to detect due to their greater effect on local gene genealogies (Figures S3–S5).

### Rescue by migrant alleles (MIG)

#### The probability of rescue and soft selective sweeps

Rescue can also arise from beneficial alleles that arrive via migration. Assuming that the number of migrant alleles that replace a resident allele each generation is Poisson with mean *m*, the waiting time until the first successful migrant is exponential with rate *λ* = *mP*_est_. Given the population is expected to persist in the absence of beneficial alleles for log(*N*(0))/*d* generations, the probability of rescue is therefore roughly the probability the first successful migrant arrives by then,

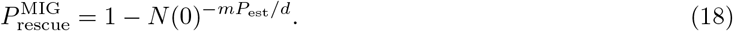

Under rescue from standing variation or new mutation we derived the probability and extent of soft selective sweeps from the forward-time process. In contrast, under rescue from migration we use the coalescent. In particular, Equation 6 shows that the per generation probability of migration and coalescence depend on population size and beneficial allele frequency in the same way. This similar form implies that the relative rates at which lineages coalesce and migrate at the selected site does not depend on the population size and allele frequency. Pennings and Hermisson (2006*a*) used this fact to show that, in an ideal population of constant size, the number of unique migrant haplotypes contributing to a present day sample, as well as their proportions, is described by Ewens’ sampling formula (pages 334*ff* in Ewens, 2004) when we replace *θ* with 2*m*. Powerfully, this results holds even in non-ideal populations of changing size (as briefly noted by Pennings and Hermisson, 2006*a*, p. 1081-1082) – including during evolutionary rescue – as long as the relationship between the effective and census population sizes remains the same (i.e., if *N*_*e*_(*t*)/*N*(*t*) is a constant; in which case we now replace *θ* with 2*mN*_*e*_/*N* in Ewens’ sampling formula). Thus the softness of a sweep from migration depends only on the migration rate and variance in gamete numbers (*σ*^2^, which determines *N*_*e*_/*N*; see Supplementary material: Genetic drift in the simulated lifecycle), and is the same during rescue as it is in a population of constant size (i.e., it is independent of *d*). The analogous result for rescue by *de novo* mutation does not hold (as it does for a population of constant size, Pennings and Hermisson, 2006*a*), since the rate of mutation is not inversely proportional to population size (Equation 6).

Here we use just two properties of Ewens’ sampling formula, the expected number of unique migrants among a sample of size *N*(page 336 in Ewens, 2004),

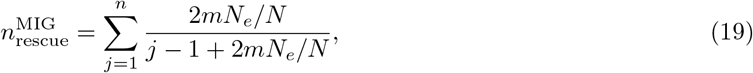

and the probability that this is more than two (equation 10.9 in Ewens, 2004),

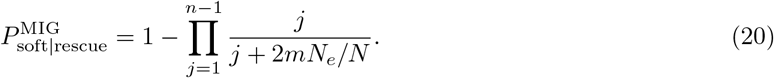

Figure 7 shows that these formulas perform very well, even when we sample the entire population (*n* = 2*N*(0)).

**Figure 7:**
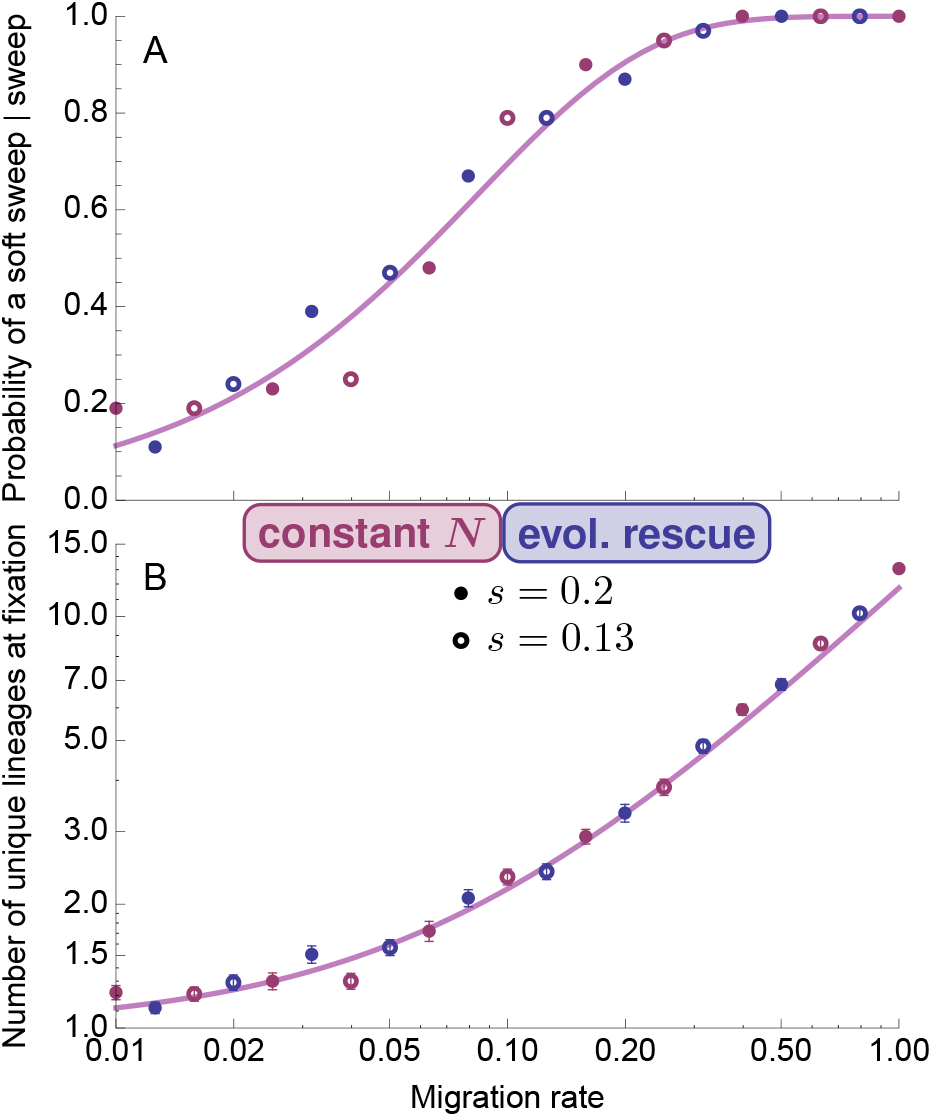
Soft sweeps from migrant alleles. (A) The probability that more than migrant lineage has descendants at the time of fixation in rescue (blue; *d* = 0.05) or in a population of roughly constant size (red; *d* = 0) as a function of the migration rate, *m*. (B) The expected number of migrant lineages that have descendants at the time of fixation. The curves are Equations 20 (panel A) and 19 (panel B) using *n* = 2*N*(0) (i.e., sampling the entire population). Each point is based on 100 replicate simulations where a sweep (and population recovery) was observed. Error bars in (B) are standard errors. Parameters: *N*(0) = 10^4^.

#### Population dynamics and the coalescent

In Supplementary material: Population dynamics and the coalescent under rescue by migration we derive the expected backward-time dynamics and structured coalescent under rescue by migration. The results are closely analogous to rescue from mutation. We do not explore genetic signatures at linked neutral sites in this case as these depend on the demographic history of the metapopulation. Previous work has explored some potential signatures of migrant sweeps in populations of constant size (e.g., Setter et al., 2019).

## Discussion

Here we have explored genetic signatures of evolutionary rescue by a selective sweep. By allowing demography to depend on the absolute fitness of the genotypes that comprise the population we explicitly invoke a feedback between demography and evolution. This feedback restricts the range of dynamics, and thus the signatures, that one should expect to observe. We find that, because the probability an allele with a given selective advantage establishes is reduced in declining populations (Equation 2; see also Otto and Whitlock, 1997), selective sweeps causing rescue are expected to be harder than those in populations of constant size when sweeps arise from standing genetic variance or recurrent mutation (Figures 1 and 6; consistent with Wilson et al., 2014 and Wilson et al., 2017). Further from the selected locus, the demographic bottleneck experienced during rescue increases the rate of coalescence relative to mutation and recombination (Figures 3 and S3), creating wider dips in relative diversity (Figures 4 and S4) and lower absolute diversity genome-wide (Figure S1; consistent with Innan and Kim, 2004). Like absolute diversity, Tajima’s *D* captures both the hardening of the sweep and the demographic bottleneck, causing *D* to often reach both higher and lower values under rescue (Figures 5 and S5). These differences between evolutionary rescue and standard sweeps all become larger under weaker selection (where the heterozygote has a smaller growth rate, *s*/2 − *d*) as the slower sweeps that result imply deeper, longer bottlenecks during rescue. If we instead compare rescue to populations bottlenecked to the same effective population size during the sweep, we find that the remaining signatures of rescue are harder sweeps that take longer to complete, allowing more time for coalescence and hence lower genome-wide diversity and elevated linkage disequilibrium (Figure S8). The subtle difference in the timing of recombination between the two scenarios, with recombination overlapping more with coalescence during rescue, may also increase variation in the number of derived mutations at a site (as expected following soft sweeps from recurrent mutation in populations of constant size; Pennings and Hermisson, 2006*b*), giving rise to site frequency spectra with perhaps slightly “flatter bottoms” (Figure S7).

In contrast to standing variance or mutation, when sweeps arise from a constant rate of migration demography has no affect on the number of beneficial alleles that establish (as briefly noted by Pennings and Hermisson, 2006*a*) and thus rescue has no affect on the hardness of the sweep (Figure 7). Further, because the rates of coalescence and migration are both inversely proportional to the number of beneficial alleles (*N*(*t*)*p*(*t*), Equation 6, Figure S10), the distribution of the number and frequency of migrant haplotypes spanning the selected site is given by Ewens’ sampling formula (Ewens, 1972), with mutation replaced by migration (Pennings and Hermisson, 2006*a*). The signatures of rescue by migration at linked neutral loci depend on the demographic and selective history of the metapopulation, which we do not explore here.

Evolutionary rescue has been explored theoretically (e.g., Gomulkiewicz and Holt, 1995; Uecker and Hermisson, 2016; Anciaux et al., 2018) and observed repeatedly in both experiments (e.g., Bell and Gonzalez, 2009; Lindsey et al., 2013; Ramsayer et al., 2013) and in host-pathogen systems in nature (e.,g., Wei et al., 1995; Feder et al., 2016). More recently, a number of studies have examined genetic data following purported evolutionary rescue in the wild, including bats altering hibernation to survive white-nose syndrome (Gignoux-Wolfsohn et al., 2018), killifish deleting receptors to tolerate pollution (Oziolor et al., 2019), and tall waterhemp evolving herbicide resistance (Kreiner et al., 2019). In the latter two cases there is strong evidence of a recent selective sweep by a very beneficial allele (in one of these cases the evidence includes reduced nucleotide diversity at the selected site; Kreiner et al., 2019). Genetic evidence for a putatively recent demographic bottleneck was presented in only one of these studies (Oziolor et al., 2019), being detected from genome-wide reductions in nucleotide diversity and increases in Tajima’s *D* relative to populations that did not experience the same environmental change. Developing a statistical model to fit our theory to such data in a more quantitative way could lead to parameter estimates, which could then inform how likely rescue may have been (or if persistence was assured regardless, *d* ≈ 0). To do so one would also have to incorporate the time since fixation as the site frequency spectra and linkage disequilibrium will begin a return to their neutral expectations following the sweep and bottleneck (e.g., the sweep signatures in Tajima’s *D* and linkage disequilibrium decay within ~ 2*N*_*e*_ generations; Przeworski, 2002). Ignoring the time since fixation would therefore cause underestimates of *s* and *d*, making rescue look more like a weaker sweep in a constant population. In contrast, the deeper and wider dips in genetic diversity observed under rescue would likely inflate estimates of selection if demography was ignored.

A lingering question that helped motivate this study is whether one could ever infer evolutionary rescue from genetic data alone. Such a possibility would greatly help assess the relevance of rescue in nature. In the empirical examples just discussed we implied that evidence for a sweep and a bottleneck is consistent with rescue. Yet stronger support for rescue would come from inferring the coincident timing of the sweep and bottleneck. Our analysis suggests that detecting coincident sweeps and bottlenecks from site frequency spectra and linkage disequilibrium will be difficult as the relative timing of coalescence and recombination leaves only subtle signatures. Consistent with this, explicit estimates of the timing of a sweep (e.g., Ormond et al., 2016) or a bottleneck (reviewed in Beichman et al., 2018) given a genetic sample collected from a single time point come with relatively large amounts of uncertainty. Sampling before and after the potential rescue event may provide one way forward, as it would help tighten the bounds on the timing of the bottleneck and sweep. However, it should be noted that even if the sweep and bottleneck appear to have co-occurred, this correlation in timing does not imply it was caused by a feedback between demography and evolution. It is, of course, difficult to say in any case – without observing replicate populations go extinct or performing experiments – whether extinction would have occurred (or will occur) without adaptive evolution, as required by the strict definition of evolutionary rescue. More experiments (such as Rêgo et al., 2019) that explore the genetic consequences of verified rescue may help develop a robust genetic signal of the feedback between demography and evolution during rescue, or at least determine if such a feedback can be detected in real genomes.

A strength of the above analysis is that we have explicitly modelled a feedback between demography and evolution, restricting the range of genetic signatures we consequently expect to observe. To take a recent example, Harris et al. (2018) have claimed that the lower reductions in genetic diversity within HIV populations adapting to less efficient drugs (as observed by Feder et al., 2016) could be due to weaker bottlenecks or slower sweeps rather than sweeps being softer, i.e., arising from multiple mutations. Fortunately, in this case genetic time-series data were available to show that the ability of HIV to reliably adapt on a short time-scale necessitates mutation rates and selection coefficients that imply adaptation by soft sweeps is likely (Feder et al., 2018). Formally modeling a feedback between demography and evolution also helps narrow the relevant parameter range. For example, under a haploid version of the model explored here (as is applicable to HIV) the minimum population size during rescue by new mutations is *N*(0)*s*(2*N*(0)*s*)^−*d/s*^/(*s − d*) (equation 22 in Orr and Unckless, 2014). Thus, for a given initial population size, *N*(0), and selection coefficient, *s*, the smallest minimum population size is *e* ln *S*/(2*s*), where *S* = 2*N*(0)*s*, implying that the smallest minimum population size consistent with the model is roughly proportional to 1/*s*. The imposed feedback between demography and evolution therefore precludes simultaneously slow sweeps and large bottlenecks (for example negating the two smallest bottleneck sizes in figure 3A of Harris et al., 2018 and constraining one to the upper right portion of figure 3A-B in Feder et al., 2018). While it is very likely in this case that soft sweeps are indeed the cause of the pattern (Feder et al., 2018), incorporating an explicit model of how demography and evolution interact could help focus future debates.

The model presented here is but one model of evolutionary rescue, which involved a number of assumptions. One of these is that the beneficial allele acts multiplicatively with the ancestral background, so that the absolute fitness its carriers, (1 + *sh*)(1 − *d*) and (1 + *s*)(1 − *d*), – and thus the probability it establishes – are affected by the decline rate of the ancestral genotype, *d*. Meanwhile, this same assumption caused the relative fitness of the mutants, 1 + *sh* and 1 + *s*, – and thus the allele frequency dynamics during the sweep, once started – to be independent of the initial decline rate (Equation 1). If, instead, we made the absolute fitness of the heterozygote and mutant homozygote independent of the initial decline rate, say 1 + *sh* and 1 + *s*, then the reverse would be true; the relative fitness of the mutant would depend on the initial rate of population decline, *d*, while its absolute fitness while establishing would not. We expect these effects would, however, largely cancel out (higher establishment probabilities will cause longer sweeps and more coalescence, but slower changes in allele frequency will allow more recombination off the sweep). In any case, because our results depend primarily on the absolute and relative fitness of the heterozygote, the alternative model just described may closely match the model analyzed in detail here when *sh* is replaced by (*sh* + *d*)/(1 − *d*).

We have also assumed that the beneficial allele acts additively with the ancestral allele at that locus (*h* = 1/2). This assumption has been made for convenience; it yields a simple explicit approximation of the allele frequency and population dynamics (Equation 5) that allows closed-form solutions of the structured coalescent (Equation 7), greatly accelerating our computation of the coalescent and pairwise diversity (Equation 9). Alternative forms of dominance are, however, likely and will impact our results. At one extreme, a completely recessive beneficial allele (*h* = 0) is much less likely to establish when compared to additivity, even in a constant population (compare Equation 2 with *v* = 1 and *ϵ* = *s*/2 to equation 15 in Kimura, 1962), making rescue nearly impossible in outcrossing populations (Uecker, 2017). In fact, any *h < d/s* will cause the heterozygote to have a negative growth rate (i.e., be subcritical), meaning that establishment will rely on the stochastic persistence of subcritical lineages until they create a supercritical mutant homozygote, analogous to the fixation of underdominant alleles or chromosomal rearrangements. As it is very unlikely that such alleles will establish in time to rescue the population (compared to supercritical heterozygotes), we have neglected this parameter range in order to focus on parameter values that are more likely to be reflected in empirical systems where rescue has occurred. It may, however, be possible to combine our approach with that taken for sweeps with arbitrary dominance in populations of constant size (Ewing et al., 2011) and the probability of establishment of underdominant alleles (e.g., equation 3 in Lande, 1979). At the other extreme, complete dominance (*h* = 1) will greatly increase the probability of establishment and rescue (Uecker, 2017), as well as population mean fitness and thus population size. All else equal, we therefore expect rescue to have less effect on the signatures of selective sweeps relative to those in populations of constant size when the rescuing allele is more dominant. Given that the marginal fitness of the beneficial allele will not depend on allele frequency under complete dominance, we expect dynamics much like the haploid model (Orr and Unckless, 2014), where simple predictions of allele frequency and population size are more accurate.

Finally, it is of course possible to model rescue under much more complex lifecycles and population structure (e.g., as expected for the evolution of malarial drug resistance; Kim et al., 2014), at least using simulations. More complex lifecycles, such as those of parasites like *Plasmodium* and HIV, could cause the bottleneck to have additional impacts on the resulting genetic signature. For example, our populations are obligate sexual out-crossers, meaning that the probability of recombination does not depend on the population size (c.f., Equation 6) as everyone must mate with some one. However, with selfing and/or facultative sex (genetic exchange), rates of recombination could be lower at lower population densities (as expected for HIV), which would increase the impact of bottlenecks on resulting genetic signatures.

Evolutionary rescue is only one example of a myriad of processes where demography and evolution feedback on one another. Our approach – combining forward-time eco-evolutionary models with coalescent theory to predict genetic signatures – could be used in many other scenarios. For instance, adaptive colonization of new habitat (a.k.a., adaptive niche expansion) is a closely related process for which a similar approach has already been taken (Kim and Gulisija, 2010). As in the case of rescue, explicitly modelling the feedback between demography and evolution in adaptive niche expansion changes the expected signatures left behind by selective sweeps as compared to (bottlenecked) Wright-Fisher populations. Such an approach is interesting from a conceptual point-of-view, improving our understanding of how eco-evolutionary dynamics affect genetic signatures. But further, given the computational power and simulation platforms available today, it is no longer necessary to restrict oneself to Wright-Fisher populations; researchers may now simulate under much more ecologically-realistic models. Developing simple approximations in parallel will become increasingly important for intuition as our models become more complex.

## Acknowledgements

We thank Pleuni Pennings and Alison Feder for helpful discussions, Ben Haller for comments on our SLiM code, and Joachim Hermisson and the anonymous reviewers for insightful comments. Financial support was provided by the Center for Population Biology at the University of California - Davis (fellowship to MMO), Banting (Canada; fellowship to MMO), and the National Institute of General Medical Sciences of the National Institutes of Health (NIH R01 GM108779, awarded to GC).

## Supplementary material: Simulated lifecycle

Here we describe the simulated lifecycle, which mimics that used in previous models of evolutionary rescue in gradually changing environments (Bürger and Lynch, 1995). Let *n*_*i*_(*t*) be the number of individuals with genotype *i* ∈ {*aa, Aa, AA*} at the beginning of generation *t*, with *N*(*t*) = ∑_*i*_ *n*_*i*_(*t*) the total population size. Viability selection, where genotype *i* survives with probability *V*_*i*_ ∈ [0, 1], is assumed to occur before reproduction. Each surviving individual then “mothers” *B* offspring, each with a randomly chosen mate (possibly oneself), and each mating produces a single offspring. If there are more than *N*(0) offspring (we assume the population starts at carrying capacity) we randomly choose *N*(0) to begin the next generation.

We next describe the deterministic allele frequency dynamics. Let *p*_*j,k*_ (*i*) be the probability a mating between genotypes *j* and *k* produces an offspring with genotype *i*. The expected number of individuals of genotype *i* after reproduction is then

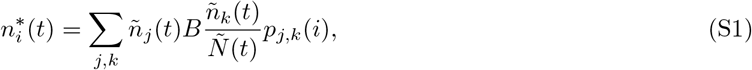

where 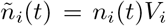 is the expected number of individuals with genotype *i* and 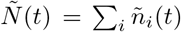 is the expected population size after viability selection. Assuming fair Mendelian transmission, the expected number of *A* alleles after reproduction is 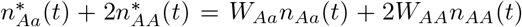, where *W*_*i*_ = *V*_i_*B* is referred to as the fitness of genotype *i*. Given that the total number of alleles after reproduction is expected to be 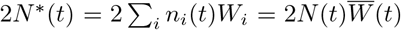, where 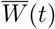 is the mean population fitness at the beginning of generation *t*, the expected frequency of allele *A* after reproduction is

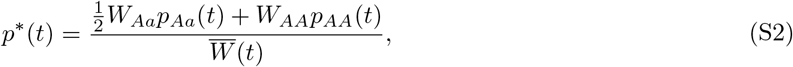

where *p*_*i*_(*t*) = *n*_*i*_(*t*)/*N*(*t*) is the frequency of genotype *i* in generation *t*. Since density-dependence is random it does not change this expectation, so that the allele frequency in next generation is *p*(*t* + 1) = *p*\*(*t*). Thus the allele frequency dynamics are the same as those in a population of constant size with relative fitnesses *W*_*i*_ (equation 5.2.3 in Crow and Kimura, 1970). Further, one can use Equation S2 to show that the genotype frequencies are expected to remain in Hardy-Weinberg proportions, allowing us to capture the dynamics of the whole system by tracking only the expected changes in the frequency of allele *A* and total population size (Equation 1).

## Supplementary material: Genetic drift in the simulated lifecycle

### Probability of establishment

In our simulated lifecycle, in generation *t* a rare allele with a viability of *V*_*i*_ survives to reproduction with probability *V*_*i*_, and given so, is present in *Y* + *Z* offspring, where *Y* is binomial with *B* trials and probability of success 1/2 (number of offspring mothered and Mendelian segregation), and *Z* is binomial with parameters *BN*(*t*) and (1/*N*(*t*))/2 (randomly chosen as a father and Mendelian segregation). With 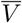 the population mean viability, the expected number of offspring produced is 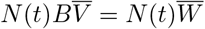. When this is less than *N*(0) there is no density dependence, so that all offspring survive to the next generation, while if 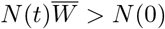 we randomly choose *N*(0) offspring to begin the next generation, implying each survives with probability 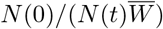.

From this we can calculate the mean and variance in the number of mutant alleles contributed to the next generation by a rare mutant allele in an individual with viability *V*_*i*_. In the absence of density dependence (e.g., in a declining population, where 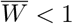) the expected number of copies of the allele contributed to the next generation is *W*_*i*_ = *BV*_*i*_ and the variance is *W*_*i*_(3 + 4(*B − W*_*i*_))/4 + *O*(1/*N*(*t*)). In a large population the heterozygote therefore has growth rate *ϵ* = *W*_*Aa*_ − 1 and variance *v* = *W*_*Aa*_(3 + 4(*B − W*_*Aa*_))/4, which we use in calculating its probability of establishment (Equation 2) for the forward-time predictions.

With density dependence (i.e., when 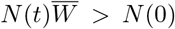) the expected number of copies contributed to the next generation is 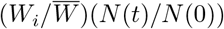 and the variance is 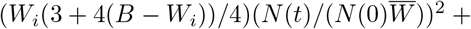 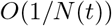. When the current population size is *N*(0) these reduce to 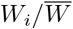 and 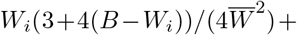 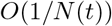. Thus the backward-time growth rate of the heterozygote in a large population of mutant homozygotes at carrying capacity is *ϵ* = 1−*W*_*Aa*_/*W*_*AA*_ and its variance is 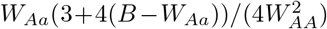, which we use in calculating the probability of establishment (Equation 2) for the effective final allele frequency (Equation 3).

### Effective population size

With slow changes in population size, *N*(*t* − 1) ≈ *N*(*t*) and the mean number of gametes contributed to the next generation by each diploid individual in the current generation is 2. The inbreeding and variance effective population size, *N*_*e*_(*t*), is then roughly (4*N*(*t*) − 2)/(*σ*^2^ + 2) (equation 7.6.4.3 in Crow and Kimura, 1970), where *σ*^2^ is the variance in the number of gametes contributed to the next generation by a parent. Therefore, in a large population, *N*(*t*) » 1, the ratio of the effective size to the census size is roughly *N*_*e*_(*t*)/*N*(*t*) ≈ 4/(2 + *σ*^2^), where *σ*^2^ depends on the particular lifecycle.

In our lifecycle, in a large population with weak selection 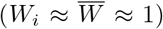 the variance is *σ*^2^ ≈ 4*B* − 3 regardless of whether or not the population is at carrying capacity (File S1). We therefore use *σ*^2^ = 4*B* − 3 to calculate the effective population size, *N*_*e*_(*t*)/*N*(*t*) ≈ 4/(2 + *σ*^2^) (see Event rates). Throughout we use *B* = 2, meaning that *σ*^2^ ≈ 5 and *N*_*e*_(*t*)/*N*(*t*) ≈ 7/4, and thus our model imparts nearly twice as much drift as a large Wright-Fisher population (where *σ*^2^ ≈ 2).

Note that drift increases with *B* because larger *B* imply that fewer individuals survive viability selection (*V* = *W/B*) but those that do have more offspring. To keep our model close to a Wright-Fisher population we use the smallest value of *B* consistent with long-term population persistence, *B* > 1. An alternative model where the expected number of offspring upon reproduction is any positive real number, say Poisson with mean *B*, would allow rates of drift closer to that of a Wright-Fisher population (by setting *B* closer to 1).

## Supplementary material: Simulation details

Forward-time simulations of the life-cycle described in Supplementary material: Simulated lifecycle were performed in `SLiM` (version 3.3; Haller and Messer, 2019) with tree-sequence recording (Haller et al., 2019).

We simulated 20 Mb chromosomes with the selected locus one of the centre bases, all other sites were neutral. We assumed a per base pair recombination rate of *r*_bp_ = 2 × 10^−8^ (i.e., 2 cM/Mb; e.g., Mackay et al., 2012) and per base mutation rate at neutral loci of *U* = 6 × 10^−9^ (e.g., Haag-Liautard et al., 2007). The recombination rate between two loci *n* bases apart was calculated as the probability of an odd number of crossover events assuming *n* independent Bernoulli trials, 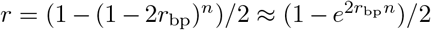 (equation 3 in Haldane, 1919), i.e., no crossover interference.

A population was considered rescued (or a sweep complete) when the beneficial mutation was fixed and the population size had recovered to *N*(0). Once a population was rescued we used `msprime` (Kelleher et al., 2016) to recapitate the population (simulate the neutral coalescent back in time from the start of the forward-time simulation, until all sites had fully coalesced) using an effective population size of *N*_*e*_(0) = 4*N*(0)/7.

From a random sample of chromosomes in the population at the time it was considered rescued, average pairwise diversity (Tajima’s *π*), Tajima’s *D*, and site-frequency spectra were calculated across 100 adjacent non-overlapping windows (i.e., each of length 200 Kb ≈ 0.4 cM) using the `diversity(), Tajimas_D()`, and `allele_frequency_spectrum()` functions in tskit (Kelleher et al., 2018). We use a sample size of 100 chromosomes throughout. Linkage disequilibrium was calculated by first identifying the segregating mutation closest to each window’s midpoint and then using `tskit.LdCalculator().r2()` to calculate disequilibrium between it and the segregating mutation closest to a recombination distance of *r* = 0.001 away.

For comparison we also run simulations with a constant expected population size by setting *d* = 0. In this case the ancestral genotype *aa* has an absolute fitness of 1, meaning that any realized population size trajectory will be a random walk with an upper boundary at *N*(0) so that extinction is assured in the long-term. However, with the parameter values used here (relatively large initial population size and fast onset of the selective sweeps), populations decline only slightly before remaining constant at the carrying capacity once the sweep has started in earnest (since the mutants have fitnesses ≥ 1). We chose to use this setup as the constant population size comparison (rather than, say, a Wright-Fisher population) because it allows us to keep the same variance in gamete numbers (affecting the probability of establishment and the rate of coalescence; see Supplementary material: Genetic drift in the simulated lifecycle) as well as the same census population size (affecting the initial and effective allele frequencies) as in the case of rescue.

## Supplementary material: Deriving the structured coalescent

Let the allele frequency and population size *τ* generations before the present be *p*′(*τ*) and *N*′(*τ*). Following Pennings and Hermisson (2006*a*), we artificially subdivide the time within a generation to be able to identify any period between two successive events in our lifecycle (Figure S6). We now go about deriving the probabilities in the structured coalescent (Equation 6).

### Migration

The number of migrant alleles that arrive each generation is Poisson with mean *m*. Given that there are 2*N*′(*τ* − 1)*p*′(*τ* − 1) beneficial alleles in the next generation, the probability that any one is a new migrant is therefore *P* = *m*/[2*N*′(*τ* − 1)*p*′(*τ* − 1)] ≈ *m*/[2*N*′(*τ*)*p*′(*τ*)], where the approximation assumes the number of beneficial alleles changes little from one generation to the next. The probability that at least one of *k* beneficial alleles is a migrant is 1 − (1 − *P*)^*k*^, which, with rare migration, is approximately

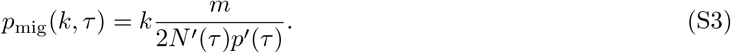

For a given probability of being replaced by a migrant allele, the rate of migration in a diploid model is half that of the haploid model (equation 15 in Pennings and Hermisson, 2006*a*, replacing *M* with *m*) as there are twice as many resident alleles.

### Mutation

The number of beneficial alleles after mutation, 2*N*′(*τ* − 4/5)*p*′(*τ* − 4/5), is the number before mutation plus the number of new mutants

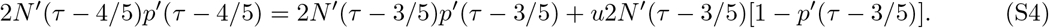

Because the population size does not change during mutation, *N*′(*τ* − 4/5) = *N*′(*τ* − 3/5), the frequency of beneficial alleles after mutation is simply

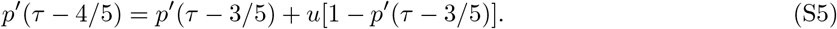

The probability a beneficial allele is a new mutant is therefore *u*[1 − *p*′(*τ* − 3/5)]/*p*′(*τ* − 4/5), which, using Equation S5, is equivalent to

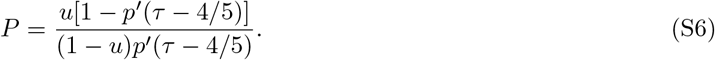

The probability that at least one of *k* beneficial alleles is a new mutant is 1 − (1 − *P*)*k*, which, when mutation is rare, is approximately *ku*[1 − *p*′(*τ* − 4/5)]/*p*′(*τ* − 4/5). With little change in allele frequency from one generation to the next this is approximately

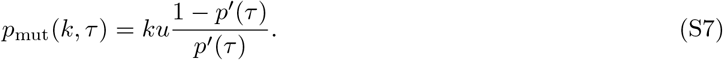

This is equivalent to the haploid result (e.g., equation 5 in Pennings and Hermisson, 2006*a*) as both the mutation rate and number of alleles are multiplied by the ploidy level, which cancels.

### Coalescence

Considering *k* beneficial alleles at the time of census, and ignoring any migration or mutation, the probability of at least one coalescence event is then

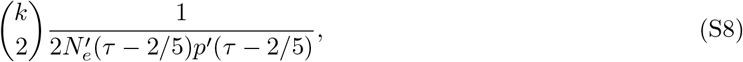

where 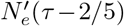 is the effective population size at the time of syngamy. When allele frequency and effective population size changes little from one generation to the next this is roughly

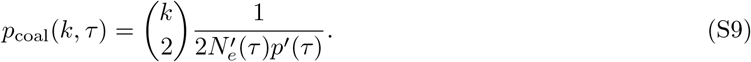

This is half the rate observed in a haploid model with the same population size (equation 5 in Pennings and Hermisson, 2006*a*) as there are twice as many alleles in a diploid population.

### Recombination

Consider a neutral locus at recombination distance *r* from the selected site. Assuming weak selection such that the survivors of viability selection remain in Hardy-Weinberg proportions, the number of alleles linked to the beneficial allele after recombination is

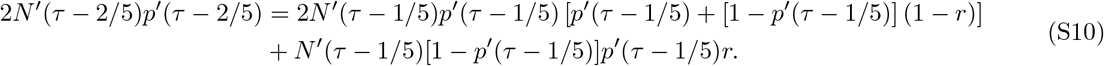

The first term on the right hand side is the number of alleles linked to the beneficial allele before recombination multiplied by the probability of being in a beneficial homozygote plus the probability of being in a heterozygote but not recombining. The second term on the right hand side is the number of alleles not linked to the beneficial allele before recombination times the probability of being in a heterozygote and recombining onto the beneficial background. The probability an allele on the beneficial background after recombination was not there before is then

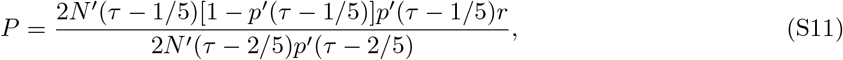

which, because recombination does not change allele frequency or population size, is

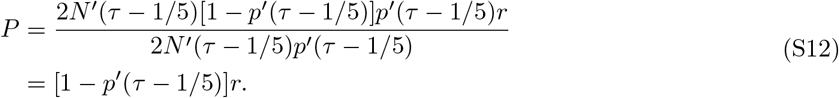

The probability at least one of *k* alleles on the beneficial background recombines off is 1 − (1 − *P*)^*k*^, which, when recombination is rare, is approximately *kr*[1 − *p*′(*τ* − 1/5)]. Assuming allele frequency changes little through one bout of selection this is *kr*[1−*p*′(*τ*)]. Finally, assuming migration, mutation, and coalescence are rare, the probability that none of *k* beneficial alleles migrate or mutate times the probability none coalesce times the probability at least one of the *k* linked alleles recombines off is roughly (table 1 in Hudson and Kaplan, 1988)

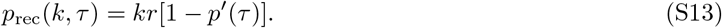

## Supplementary material: A simultaneous, but independent, bottleneck and sweep from SGV

In Rescue from standing genetic variation (SGV) we have compared the genetic signatures of rescue to those from sweeps in populations of constant size. Two of key differences we have identified, lower absolute diversity and positive Tajima’s *D* at unlinked sites under rescue, arise simply because of the population bottleneck. This suggests that it will be yet harder to distinguish rescue from a sweep that occurs during a bottleneck, where the sweep and bottleneck are coincident but independent. Taking this argument to its logical limit, if the population size dynamics during an externally forced bottleneck are exactly like those expected under rescue, then the expected genetic signatures will be identical. As a middle-ground, we can compare an instance of rescue that has an expected effective population size of *N*_*e*_ during its sweep to a population that is bottlenecked to a constant effective population size *N*_*e*_ during its sweep (let’s call the latter the ‘coincident’ scenario). Then, under the same allele frequency dynamics (which will not be exactly true as population size and decline rate affect the effective initial and final allele frequencies), both populations would have similar genome-wide diversity and Tajima’s *D* values.

In File S1 we show that coincident scenario produces patterns of relative diversity at the selected site like sweeps in populations of constant size (i.e., like the red curve at 0cM in Figure 4), meaning that rescue causes deeper dips in diversity than a coincident sweep and bottleneck. This is because the coincident scenario assumes smaller population sizes at the beginning and end of the sweep, causing higher effective initial frequencies and lower effective final frequencies, which shortens the length of the sweep and thus increases the probability that no events occur in the history of the sample during the sweep, *P*_*∅*_(*k, t*_*f*_). These shorter sweeps also mean that the coincident scenario leaves less time for coalescence on the ancestral background and thus produces higher genome-wide absolute diversity than populations that have been rescued. In contrast, rescue and the coincident scenario similarly slow the recovery of relative diversity as we move away from the selected site, causing their dips in diversity to have similar width (i.e, like the blue curve in Figure 4). This is because both scenarios increase rates of coalescence early in the sweep, preventing recombination off and thus lowering *P*_off_(*k, t*_*f*_). The shape of Tajima’s *D* values across the genome are much the same in the coincident scenario as they are after a sweep in a population of constant size (i.e., like the red curves in Figure 5), just shifted up so that the background levels closely match those under rescue (i.e., they have intercepts like the blue curves in Figure 5). In summary: for a given strength of selection and effective population size during a sweep, the dynamics of rescue produce 1) deeper dips in diversity, 2) lower absolute diversity genome-wide, and 3) a greater range of Tajima’s *D* values. These differences arise because the lower population sizes at the beginning and end of the coincident scenario cause faster establishment and fixation, leading to shorter sweeps.

To explore whether there is additional signal to be gained from the different timings of coalescence and recombination in the coincident and rescue scenarios we have also looked at the full site frequency spectra arising from simulations (of which pairwise diversity and Tajima’s *D* are summaries; Wakeley, 2009, p. 116). Figure S7 shows the site frequency spectra at three recombination distances for a case where relative diversity and Tajima’s *D* values are very similar in the two scenarios (*s* = 0.2, *κ* = 10; see File S1). Far from the selected site the entire spectra are very similar in the two scenarios, both documenting a recent short drop in population size (a deficiency of rare alleles, as indicated by positive genome-wide Tajima’s *D*). As we move towards the selected site the patterns diverge slightly, the harder sweep during rescue now causing a larger deficit of intermediate frequency alleles and a larger excess of high frequency alleles (as indicated by a slightly more negative Tajima’s *D* at the shoulders of the selected site). At selected site the spectra are again very similar, although with more noise as there are less polymorphic sites. Thus the strongest signal of rescue vs. a bottlenecked sweep seems to be provided at moderately linked sites, where the relative timings and strengths of coalescence and recombination have the most impact. For instance, the timing and probability of coalescence at 1cM is nearly identical in rescue vs. the coincident scenario here, but the probability of recombination is slightly reduced and tends to occur slightly later under rescue. This reduction in recombination is predicted to create phylogenies that are more star-like, which may explain much of the greater deficit of intermediate frequency mutations. The fact that recombination is also delayed and thus overlaps more with coalescence, in rescue relative to the coincident scenario, may help explain the perhaps flatter spectrum across low to high frequency mutations near the selected site, as the overlap creates more variance in the number of coalescence events that occur prior to the lineage recombining off the sweep (similar to the effect of recurrent mutation; Pennings and Hermisson, 2006*b*). Note that if we instead compared site frequency spectra in windows with similar absolute genetic diversity (as in figure 6 of Kim and Gulisija, 2010), differences between the three scenarios would be more apparent. Thus a combination of summary statistics may help differentiate rescue from a simultaneous, but independent, bottleneck and sweep.

Site frequency spectra (and summaries of them) are characteristics of individual loci. As a final potential signature of rescue vs. the coincident scenario we look at linkage disequilibrium, which captures correlations in the pairwise coalescence times between two loci (Wakeley, 2009, p. 236). Figure S8 shows linkage disequilibrium between neutral loci that are a recombination distance of *r* = 0.001 apart, as a function of their distance from the selected site. As we can see, the sweep causes elevated linkage disequilibrium near the selected site, as expected for sites that were segregating before the sweep (Kim and Nielsen, 2004; McVean, 2007). Meanwhile population bottlenecks tend to increase linkage disequilibrium genome wide, as expected given that bottlenecks decrease mean coalescence times more than they decrease the variance (McVean, 2002). The differences in linkage disequilibrium patterns between the rescue and coincident scenario are much greater than the differences observed in relative diversity, Tajima’s *D*, or the full site frequency spectrum. The differences in linkage disequilibrium do, however, largely mirror those observed in absolute diversity (see File S1), which can largely be explained by the shorter sweeps, and thus less coalescence, in the coincident scenario.

## Supplementary material: Population dynamics and the coalescent under rescue by migration

### Effective initial allele frequency and the backward-time dynamics

Dividing the (truncated) exponential waiting time distribution for the first successful migrant allele by the probability of rescue gives the waiting time distribution conditioned on rescue. Following the same approach as above (see Rescue by *de novo* mutation (DNM)), the effective initial frequency of the beneficial allele given rescue is then

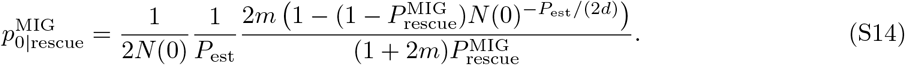

When the migration rate is small this last factor is nearly independent of *m* (analogous to the mutation case). As in the case of *de novo* mutation, to characterize the backward-time dynamics in a population of constant size we do not need to know when the sweep begins, just the effective initial frequency at this time, 1/(2*N*(0)*P*_est_).

Figure S9 compares our analytical approximations (Equations 5 and S14) against individual-based simulations. We see the predictions do fairly well for larger values of *m*, but can fair quite poorly with small *m*. In the latter case the first successful migrant allele sometimes arrives when the population is so small that the beneficial allele increases in frequency much faster than the deterministic expectation. Here we enter a different regime, which we do not attempt to approximate.

### The structured coalescent

Figure S10 shows the timing of migration relative to recombination and coalescence (Equation 7). As with rescue from standing genetic variance or mutation (Figures 3 and S3), the bottleneck increases the overall coalescence rate and shifts its timing closer to fixation, overlapping more with recombination. Migration scales with coalescence (Equation 6) and is thus similarly increased and shifted. When migration rates are small enough that the migrant allele tends to fix when the population is very small the simulations show an initial spike in expected coalescence, which is expected to greatly reduce diversity.

### Genetic signatures at linked neutral loci

The genetic signatures at linked neutral loci produced by a sweep that arises from migration depends on the history of the metapopulation, e.g., how and when the sweeps occurred in each patch and the historical migration rates between them. We therefore omit an exploration of the signatures produced in this scenario, which deserves a full and careful treatment on its own. Previous work has explored some potential signatures of migrant sweeps in populations of constant size. For example, if the historic migration rate between the migrant and focal population is low then we should expect few migrant alleles away from the selected site. Then, under low contemporary migration rates we expect a relatively hard sweep and the so-called “volcano” pattern of genetic diversity (Setter et al., 2019), where diversity is maximized at an intermediate distance from the selected site due to a more balanced presence of both migrant and non-migrant alleles.

## Supplementary figures

**Figure S1:**
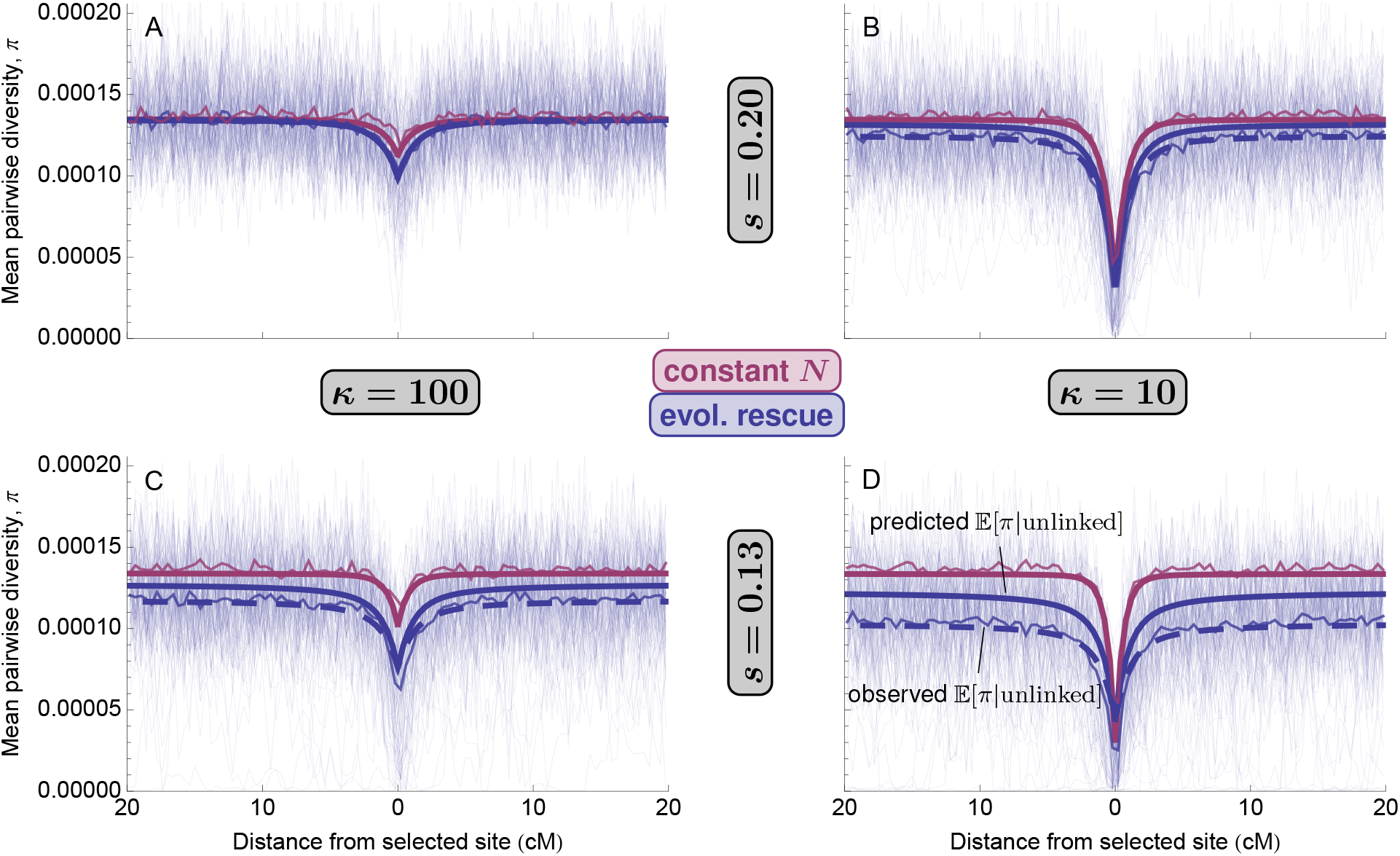
Mean pairwise diversity, *π*, after a selective sweep from standing genetic variation in evolutionary rescue (blue; *d* = 0.05) or in a population of roughly constant size (red; *d* = 0). The thick solid curves are our approximations (Equation 9). The dashed blue curves replace 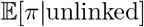 with the observed genome-wide average *π* (excluding sites within 5cM of the selected site as we simulate only a portion of a genome). The thinnest curves are 100 replicate simulations (rescue only for clarity) and the slightly thicker curves are simulation means (often obscured by prediction). Parameters: *N*(0) = 10^4^.

**Figure S2:**
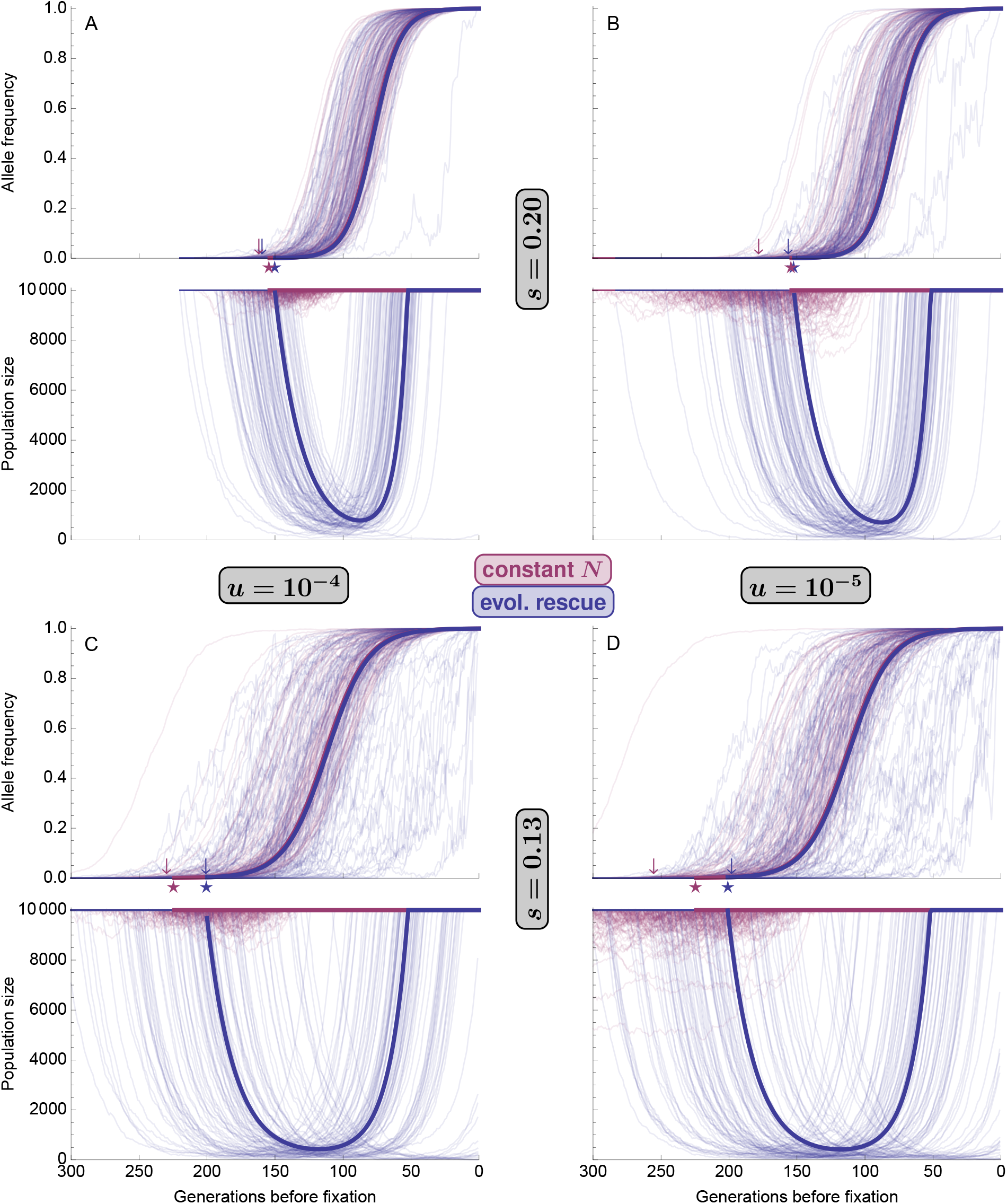
Allele frequency, *p*′(*τ*), and population size, *N*′(*τ*), at generation *τ* before fixation during a selective sweep from *de novo* mutation in evolutionary rescue (blue; *d* = 0.05) and in a population of roughly constant size (red; *d* = 0). The thick solid curves are analytic approximations (Equation 5), using 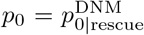 (Equation 17) as the initial frequency and *p*_*f*_ (Equation 3) as the final allele frequency. The thin curves are 100 replicate simulations where a sweep (and population recovery) was observed. The stars show the predicted time to fixation (Equation 4). The arrows show the mean time to fixation observed in simulations.

**Figure S3:**
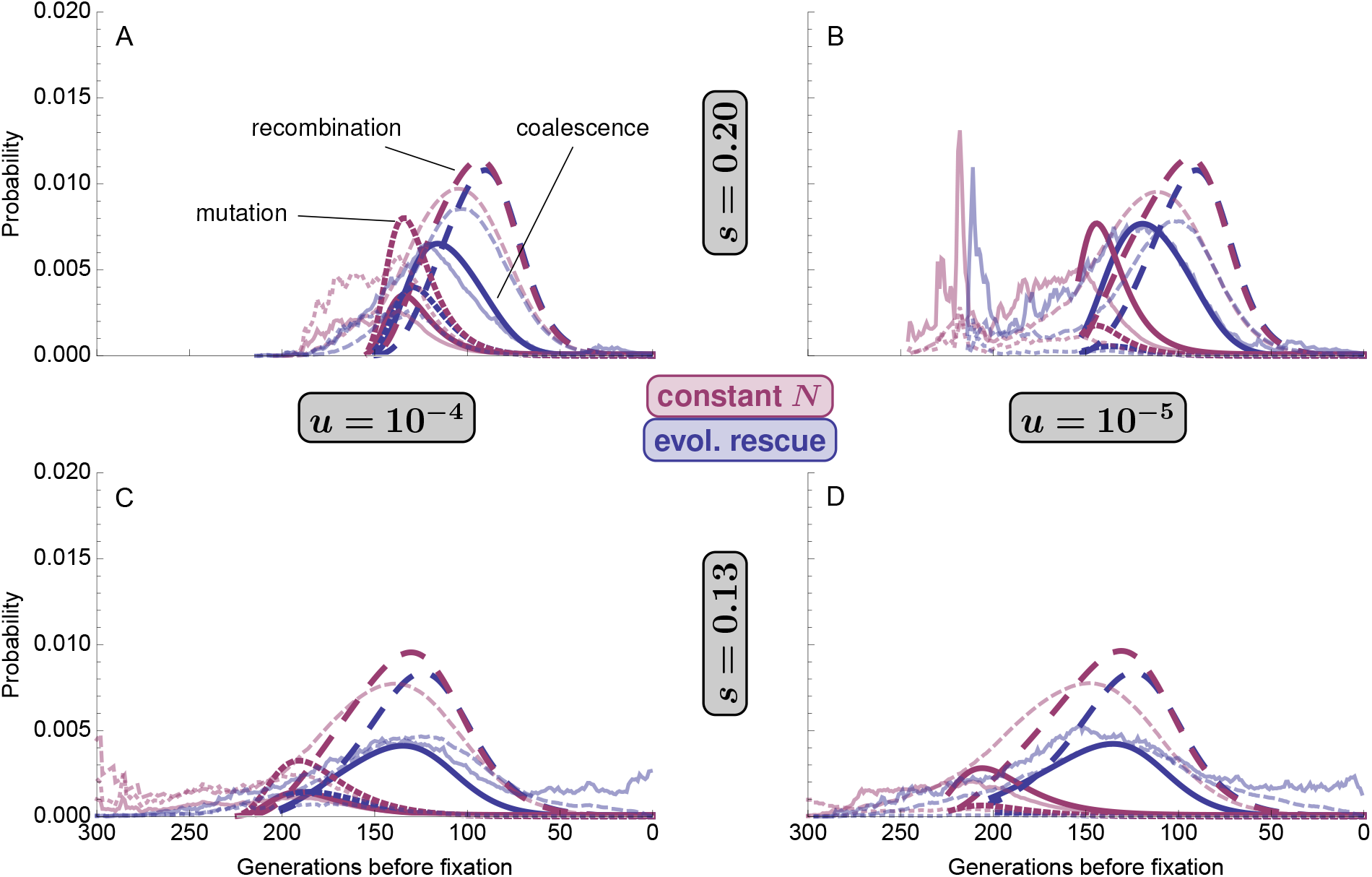
The timing of events in the structured coalescent (Equation 7) for a sample of size *k* = 2 at a linked neutral locus (*r* = 0.01) during a selective sweep from *de novo* mutation in evolutionary rescue (blue; *d* = 0.05) or in a population of roughly constant size (red; *d* = 0). Thicker opaque curves use the analytic expressions for allele frequency and population size (Equation 5) while the thinner transparent curves show the mean probabilities given the observed allele frequency and population size dynamics in 100 replicate simulations (these become more variable in the past as less replicates remain polymorphic).

**Figure S4:**
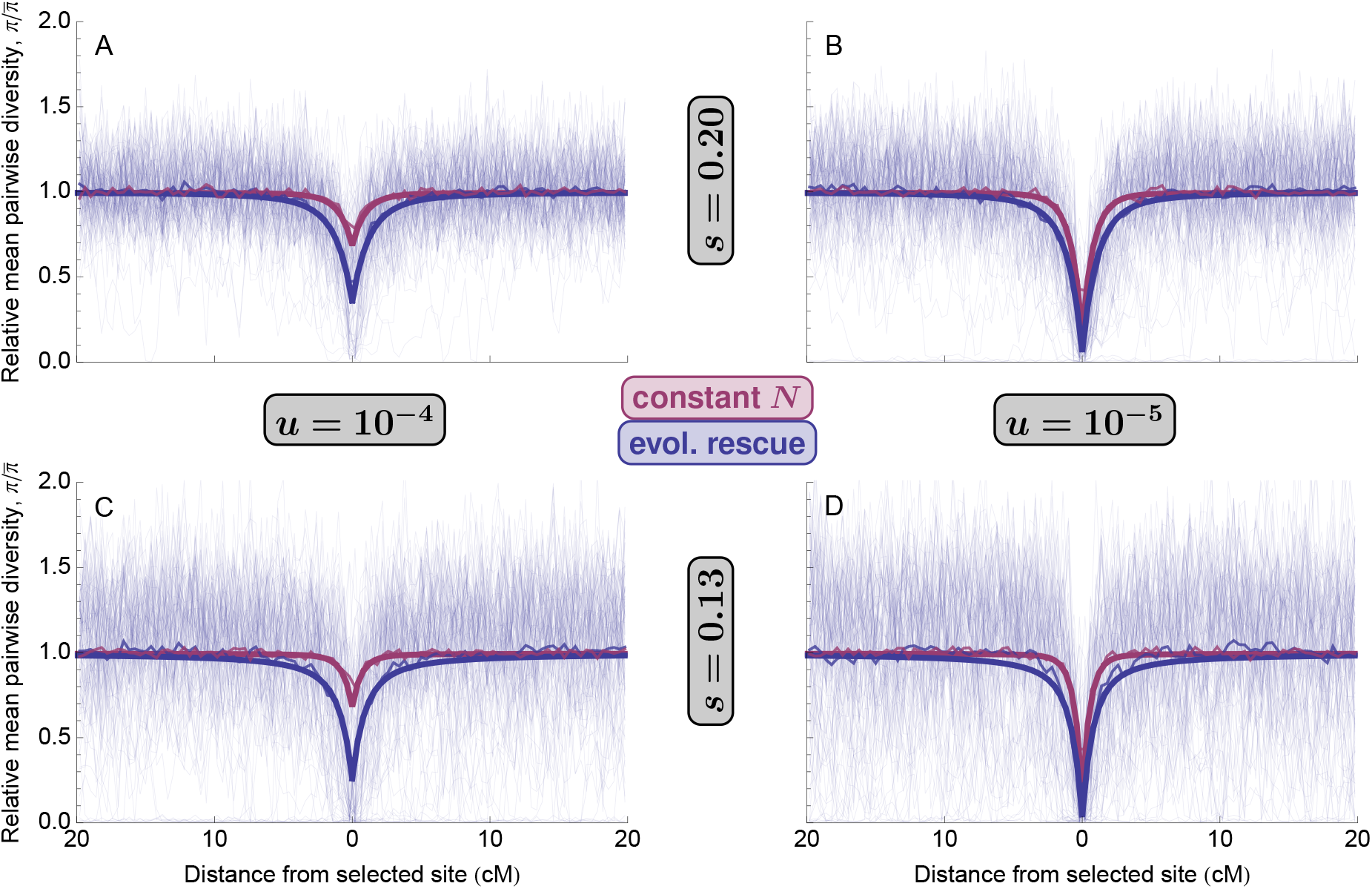
Relative mean pairwise diversity 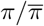, after a selective sweep from *de novo* mutation during evolutionary rescue (blue; *d* = 0.05) or in a population of roughly constant size (red; *d* = 0). The thickest curves are 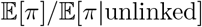 (using Equation 9). The thinnest curves are 100 replicate simulations (rescue only for clarity) and the slightly thicker curves are simulation means (often obscured by prediction). Parameters: *N*(0) = 10^4^.

**Figure S5:**
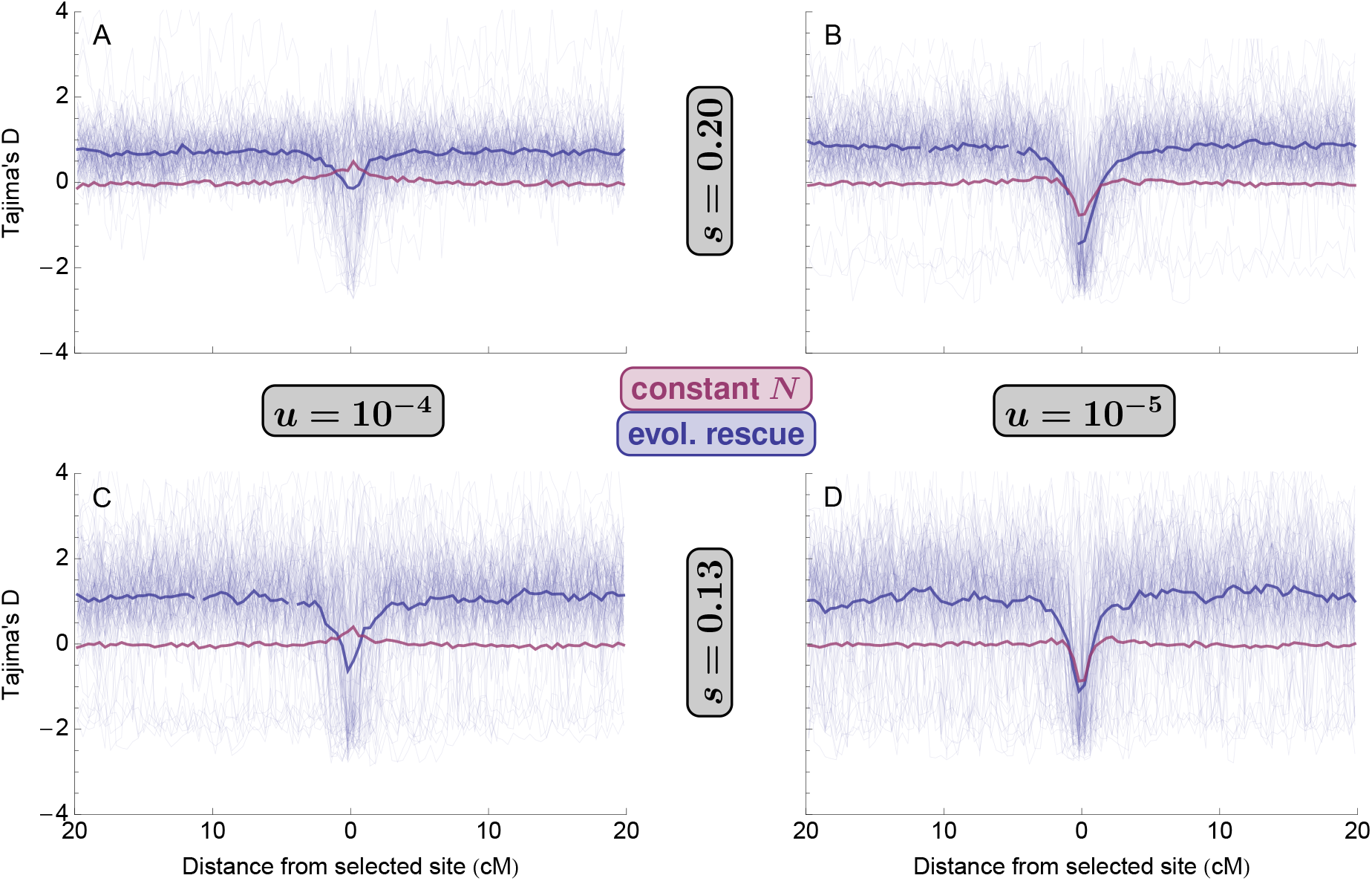
Tajima’s *D* after a selective sweep from *de novo* mutation during evolutionary rescue (blue; *d* = 0.05) or in a population of roughly constant size (red; *d* = 0). Thin curves show 100 replicate simulations (rescue only for clarity) and thicker curves show simulation means. Parameters: *N*(0) = 10^4^.

**Figure S6:**
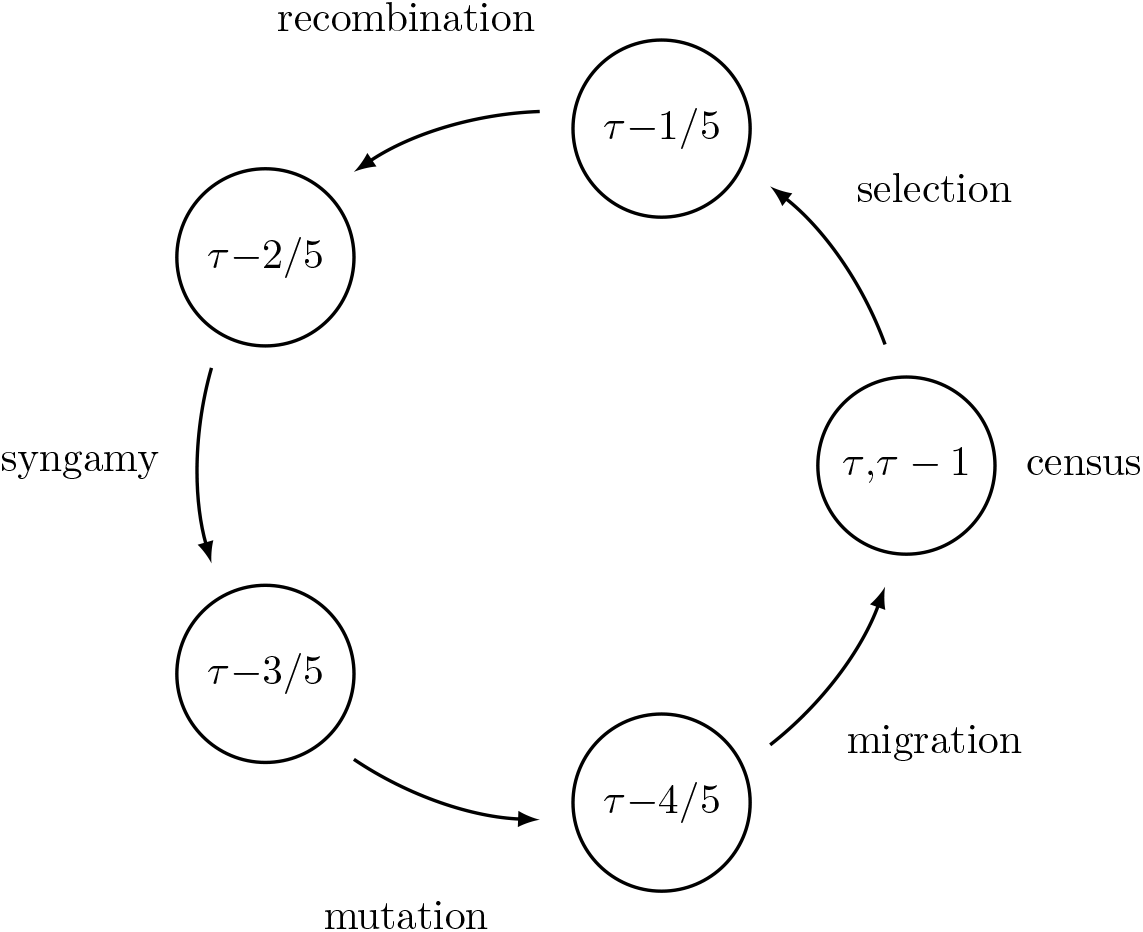
Life-cycle and time notation. The arrows indicate the forward-time direction and the numbers indicate the time before fixation (i.e., starting in generation *τ* and moving forward in time through *τ* − 1/5, *τ* − 2/5, … we arrive at generation *τ* − 1, one generation closer to fixation).

**Figure S7:**
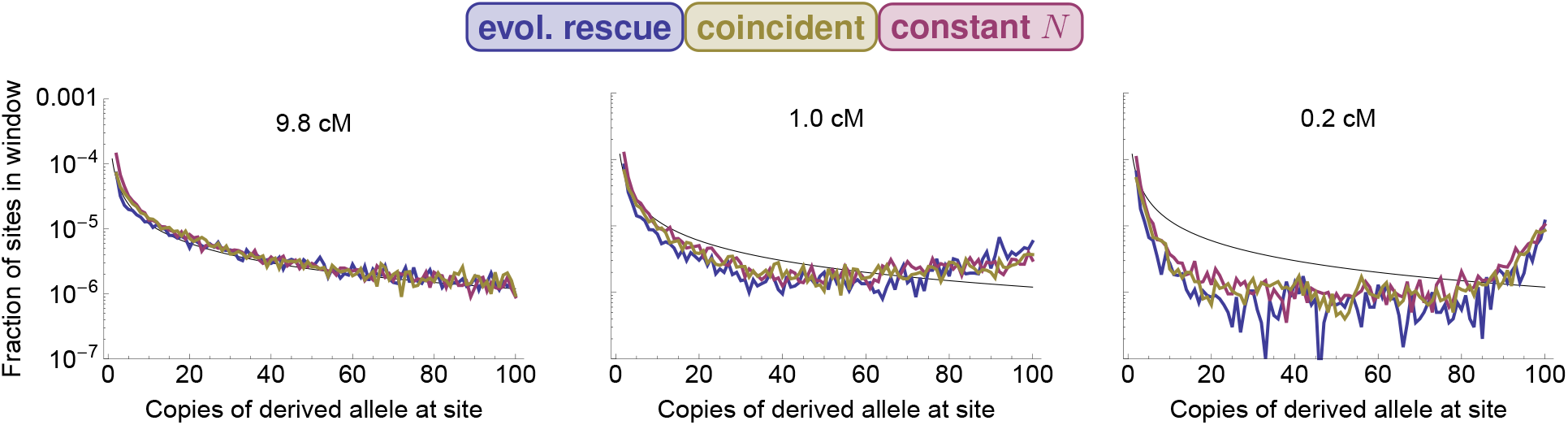
Site frequency spectra at three distances from the selected site after a selective sweep from standing genetic variation in evolutionary rescue (blue; *N*(0) = 10^4^, *d* = 0.05), in a bottlenecked population with short-term effective population size similar to that in rescue (yellow; *N*(0) = 2945, *d* = 0), or in a population of constant size (red; *N*(0) = 10^4^, *d* = 0). Blue, yellow, and red curves are mean values from 100 replicate simulations. The black curve is the neutral expectation, 4*N*_*e*_*U/x*, where *N*_*e*_ is the long-term effective population size (here 10^4^ × 4/7) and *x* is the number of copies of the derived allele. Parameters:*s* = 0.2, *κ* = 10.

**Figure S8:**
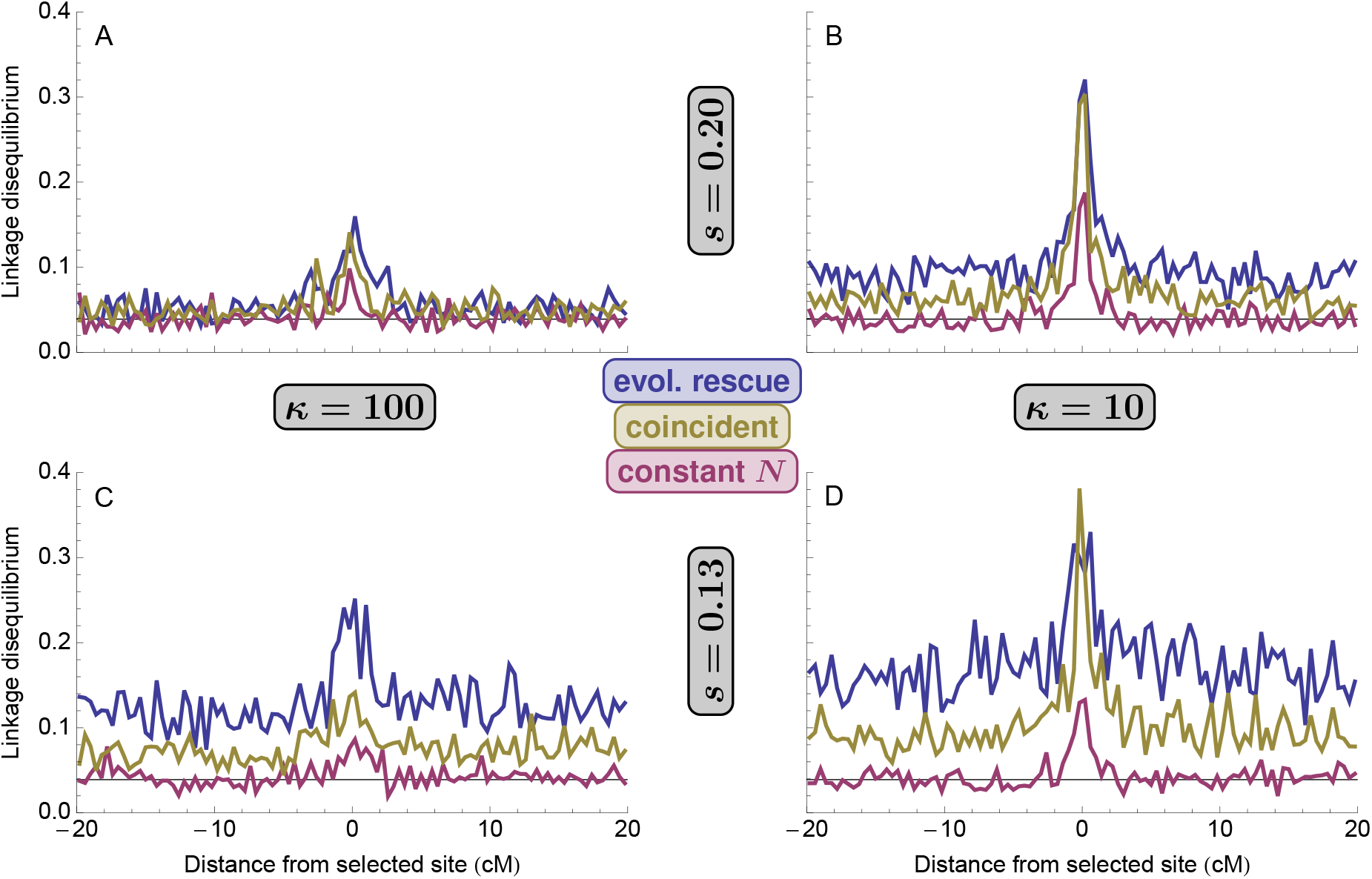
Linkage disequilibrium, *r*^2^, between neutral loci that are recombination distance 0.001 apart after a selective sweep from standing genetic variation in evolutionary rescue (blue; *N*(0) = 10^4^, *d* = 0.05), in a bottlenecked population with short-term effective population size similar to that in rescue (yellow; *N*(0) = 2945, *d* = 0), or in a population of constant size (red; *N*(0) = 10^4^, *d* = 0). The black line is the neutral expectation (equation 7.31 in Wakeley, 2009).

**Figure S9:**
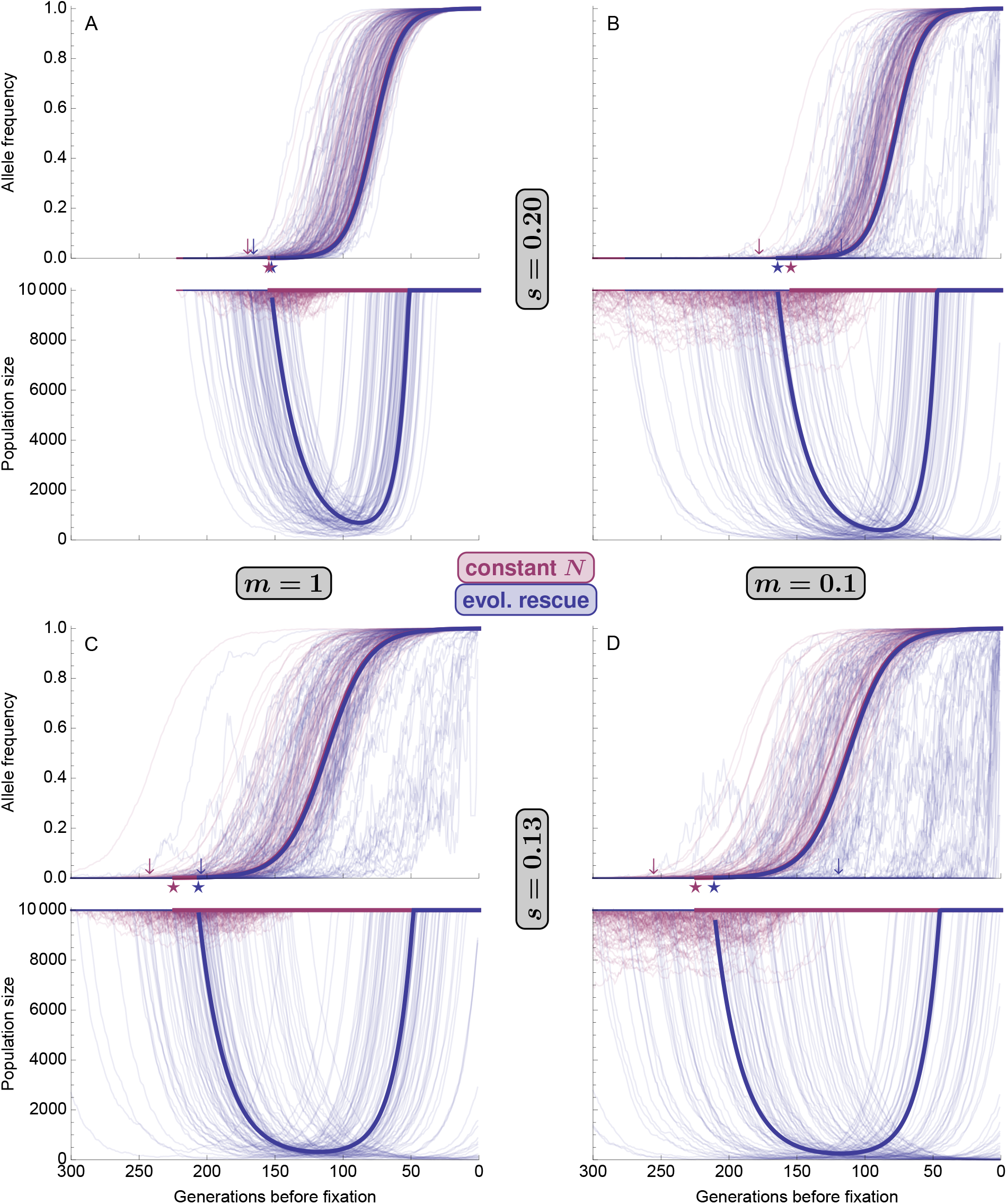
Allele frequency and population size during a selective sweep from migration in evolutionary rescue (blue; *d* = 0.05) or in a population of roughly constant size (red; *d* = 0). The thick solid curves are analytic approximations (Equation 5), using 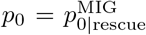 (Equation S14) as the initial frequency and *p*_*f*_ (Equation 3) as the final allele frequency. The thin curves are 100 replicate simulations where a sweep (and population recovery) was observed. The stars show the predicted time to fixation (Equation 4). The arrows show the mean time to fixation observed in simulations.

**Figure S10:**
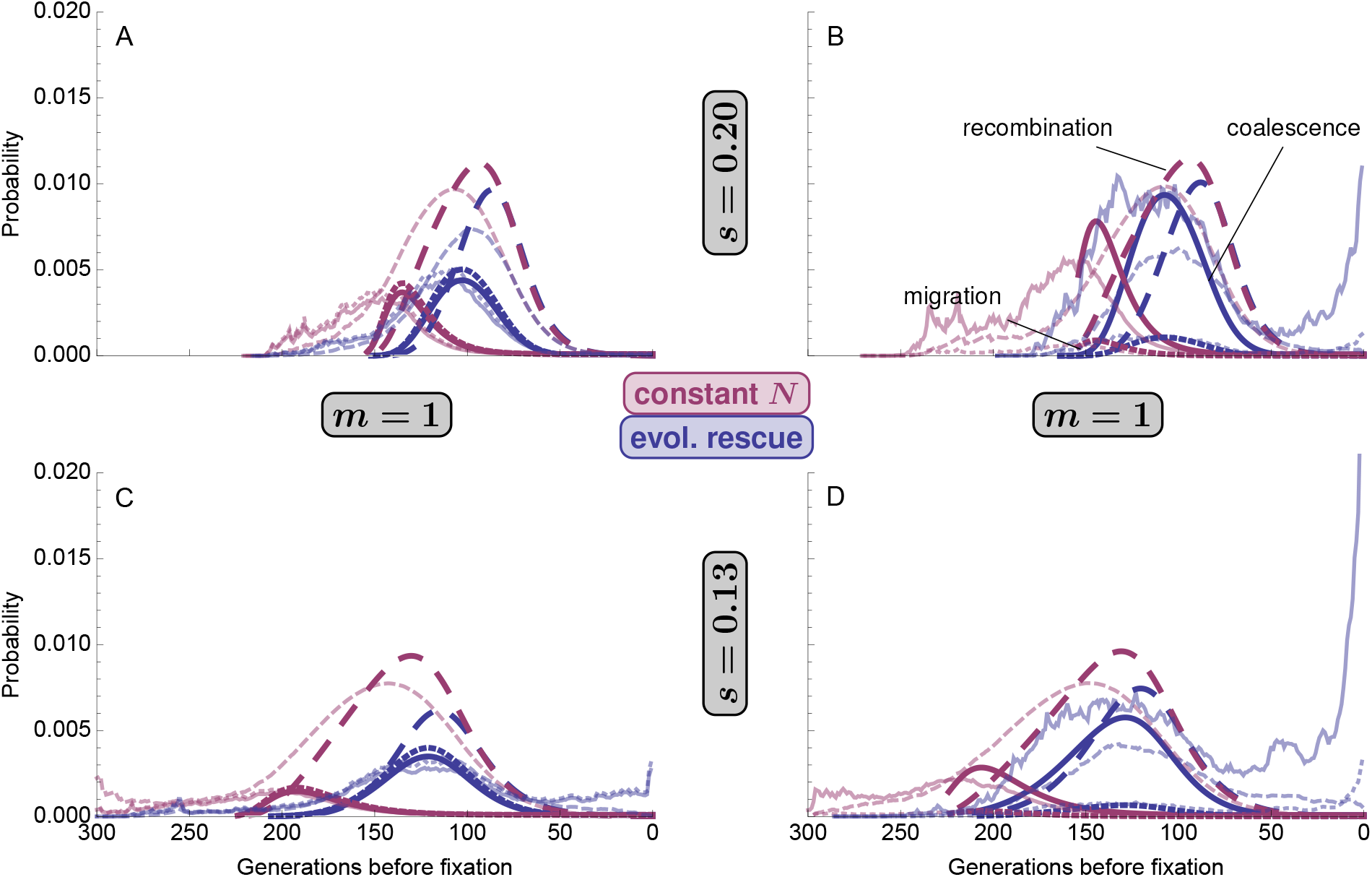
The timing of events in the structured coalescent (Equation 7) for a sample of size *k* = 2 at a linked neutral locus (*r* = 0.01) during a selective sweep from migration in evolutionary rescue (blue; *d* = 0.05) or in a population of roughly constant size (red; *d* = 0). Thicker opaque curves use the analytic expressions for allele frequency and population size (Equation 5) while the thinner transparent curves show the mean probabilities given the observed allele frequency and population size dynamics in 100 replicate simulations (these become more variable in the past as less replicates remain polymorphic).

